# MND1 enables repair of two-ended DNA double-strand breaks

**DOI:** 10.1101/2022.08.24.505076

**Authors:** Lisa Koob, Anoek Friskes, Louise van Bergen, Femke M. Feringa, Bram van den Broek, Emma S. Koeleman, Michael Schubert, Vincent A. Blomen, Thijn R. Brummelkamp, Lenno Krenning, René H. Medema

## Abstract

Faithful and timely repair of DNA double-strand breaks (DSBs) is fundamental for the maintenance of genomic integrity. Here, we demonstrate that the meiotic recombination co-factor MND1 facilitates the repair of DSBs in somatic cells. We show that MND1 localizes to DSBs, where it stimulates DNA repair through homologous recombination (HR). Importantly, MND1 is not involved in the response to replication-associated DSBs, implying that it is dispensable for HR-mediated repair of one-ended DSBs. Instead, we find that MND1 specifically plays a role in the response to two-ended DSBs that are induced by IR or various chemotherapeutic drugs, specifically in G2. MND1 localization to DSBs is dependent on resection of the DNA ends, and seemingly occurs through direct binding of MND1 to RAD51-coated ssDNA. Importantly, the lack of MND1-driven HR repair directly potentiates the toxicity of IR-induced damage, which could open new possibilities for therapeutic intervention, specifically in HR-proficient tumors.

## Introduction

The integrity of the human genome is constantly challenged by a wide variety of endogenous and exogenous DNA damaging sources. To resolve DNA breaks, cells have evolved an intricate network of proteins that sense and repair the damage, collectively known as the DNA damage response (DDR) (Blackford and Jackson, 2017; Ciccia and Elledge, 2010; Jackson and Bartek, 2009). A dysfunctional DDR has been shown to drive cancer progression and evolution by accelerating the accumulation of mutations (Hanahan and Weinberg, 2011; Tubbs and Nussenzweig, 2017). Hence, mutations in DDR genes are commonly found as driver mutations in cancer. Such mutations often lead to inactivation of specific repair pathways and, as a consequence, cancers driven by mutations in DDR genes are often hypersensitive to loss or chemical inhibition of the alternative repair pathways (Marshall and Antonarakis, 2020; O’Connor, 2015; Pilié et al., 2019). This indicates the importance of a careful delineation of distinct repair pathways, as their redundancies create the potential for novel anti-cancer therapies.

A widely effective DNA damage-inducing cancer treatment in the clinic is radiotherapy (Huang and Zhou, 2020). The detrimental effect of radiotherapy is mainly based on the induction of DNA double-strand breaks (DSBs) by ionizing radiation (y-irradiation, IR). DSBs are one of the most deleterious types of DNA damage (Nickolo et al., 2020; Toulany, 2019). DSBs are repaired by a plethora of different repair pathways: non-homologous end-joining (NHEJ), homologous recombination (HR), single strand annealing (SSA), or microhomology-mediated end-joining (MMEJ). While NHEJ functions throughout the cell cycle, HR and SSA are mainly restricted to S and G2 phases (Hustedt and Durocher, 2017), whereas MMEJ requires passage through mitosis (Llorens-Agost et al., 2021; Wang et al., 2018). This is in part because HR is dependent on a homologous DNA strand, which is only present post-replication during S phase. For HR to occur, DNA resection is required to generate a template for strand invasion. In contrast, NHEJ only requires minimal DNA end processing before ligation of broken ends (Chang et al., 2017; Chatterjee and Walker, 2017). Resection generates stretches of single-stranded DNA (ssDNA) that are coated by RPA. Later, RPA is exchanged for the RAD51 recombinase, which enables homology search and strand invasion of the sister chromatid (Jensen et al., 2010; Sullivan and Bernstein, 2018). RAD51-mediated homology search and strand invasion is facilitated by its ATPase enzymatic function, whose inhibition abolishes DNA strand exchange and successful HR repair (Bonilla et al., 2020; Huang and Mazin, 2014).

Although much is known about the DDR network, genome-wide screening in the context of various DNA damaging treatments continues to identify genes that are involved in DNA repair. This is highlighted by the recent discovery of DDR factors like Shieldin, ELOF1 and ERCC6L2 (Dev et al., 2018; Francica et al., 2020; Noordermeer et al., 2018; Olivieri et al., 2020). Further identification of DDR genes will generate a more complete picture of the distinct repair pathways, potentially leading to the identification of novel (adjuvant) therapeutic targets. Therefore, we set out to identify factors that limit the toxicity of DSBs that are induced by IR. For this, we performed a haploid genetic screen aimed to identify genes involved in the cellular survival in response to IR. The screen identified many genes with well-established roles in the DNA damage response, as well as several genes with no previously described role in DNA repair.

One of the most prominent hits in our screen was MND1, loss of which resulted in a marked increase in sensitivity to IR. This finding was surprising as MND1 is known for its role in homology search and recombination during meiosis, but not in somatic cells. Specifically, loss of MND1 has been shown to cause persistence of meiotic DSBs and results in the formation of non-homologous synapses (Leu et al., 1998). During meiotic recombination, MND1 binds its co-factor HOP2 and stabilizes both RAD51- and DMC1-coated presynaptic filaments, which facilitates strand invasion and D-loop formation (Bugreev et al., 2014; Chi et al., 2007; Pezza et al., 2007). MND1 has not been directly implicated in DSB repair during the somatic cell cycle. However, MND1 has been shown to facilitate the alternative lengthening of telomeres (ALT) (Cho et al., 2014), a process of telomere replication that is dependent on many established HR factors.

Here we describe that MND1 facilitates DSB repair through HR also during the somatic cell cycle. We find that loss of MND1-HOP2 complex sensitizes cells to DSBs induced by IR and various chemotherapeutic drugs. Interestingly, we find that MND1 is dispensable for HR-dependent repair of replication-associated breaks, indicating that targeting MND1 can be a way to inhibit some, but not all, HR-dependent repair. MND1 localizes readily to DSBs where it facilitates the timely resolution of RAD51 foci and stimulates HR. Consequently, MND1 loss potentiates the G2 DNA damage checkpoint, causing hypersensitivity to DNA damage during G2 phase. Therefore, we conclude that MND1 has a critical role in the repair of DSBs via HR during the somatic cell cycle.

## Results

### A haploid genetic screen identifies MND1 loss as sensitizing towards y-irradiation

To identify novel factors involved in the DNA damage response (DDR), we performed a genetic perturbation screen using random mutagenesis by gene-trap insertion in haploid human cells (HAP1 cells) (Blomen et al., 2015)). Gene function is mainly disrupted upon integration of the gene-trap in the sense orientation. As such, gene essentiality is determined by calculating the ratio of sense versus antisense integrations. In this study, mutagenized cells were treated with 1Gy IR five times every other day, and compared to untreated control (WT – 0Gy) cell populations (Blomen et al., 2015) (Figure 1A, one representative WT dataset is depicted). After 10 days of culture, cells were collected and gene-trap insertions were determined by next-generation sequencing (see methods for a detailed description of the screen setup). We defined a gene as a hit when it is was enriched for antisense-orientation gene-trap insertions in the irradiated cell population (odds ratio ≤0.8), and the insertion ratio was significantly different compared to the four independent control datasets (p-value ≤0.05) (Blomen et al., 2015)). This resulted in a list of 261 genes (Suppl. Table 1, highlighted in brown in Figure 1B). Gene set enrichment analysis of the screen hits identified multiple pathways involved in the DNA damage recognition and repair to be significantly enriched (Figure 1C), indicating that our screen setup is able to identify DDR factors.

**Figure 1:**
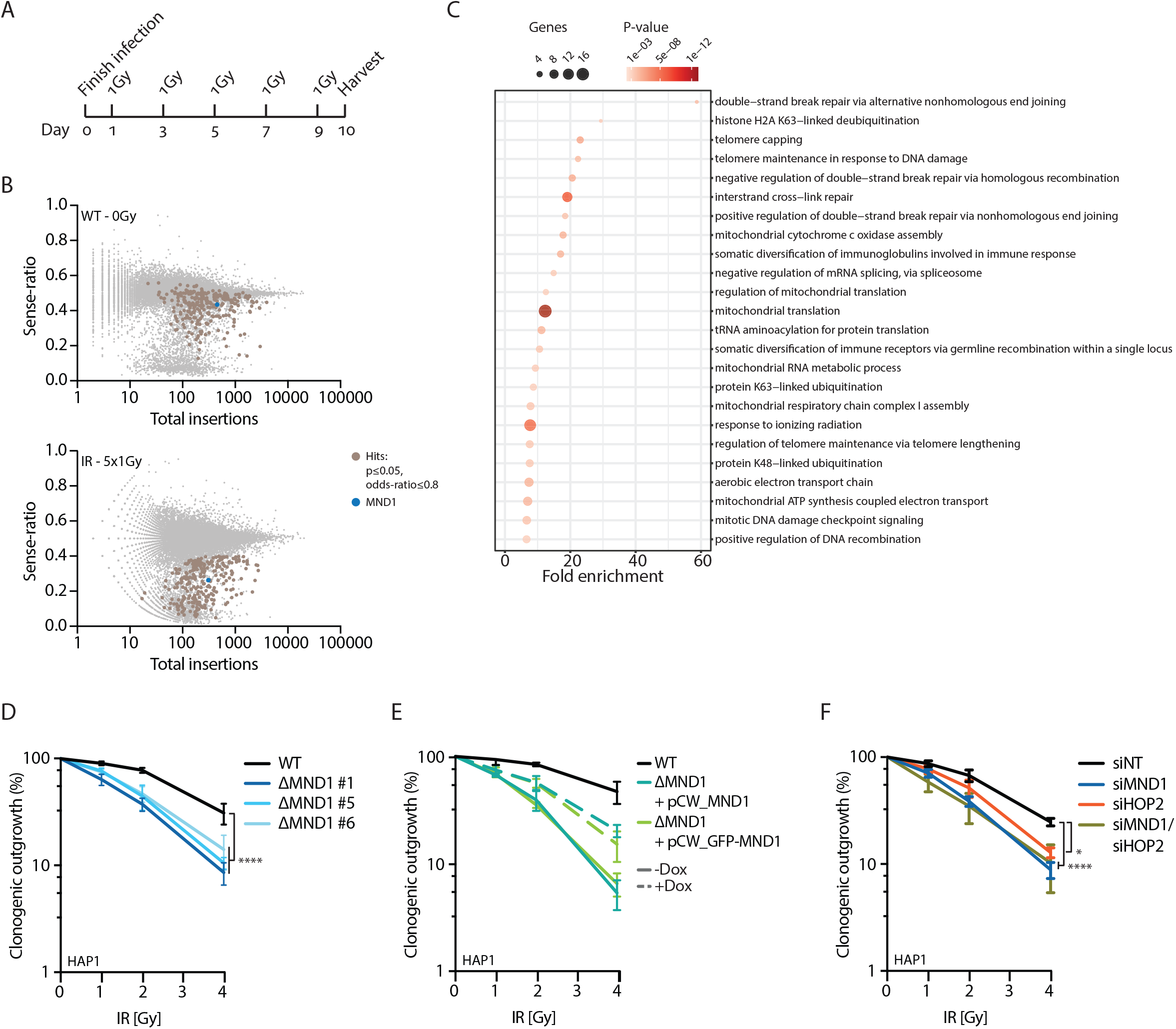
Haploid genetic screen identifies MND1 as a sensitizer to IR-induced DNA damage. A) Treatment schedule of the haploid genetic screen. Cells were treated with 1Gy of IR every other day, five times. The cells were harvested 10 days after the start of the screen. B) Results from the HAP1 genetic screen (Control – 0Gy (one dataset is displayed) and IR – 5×1Gy) are shown by plotting the relative amount of mapped sense integrations over total insertions. Hits are defined with p-values ≤0.05 when compared against all four WT datasets and an odds-ratio ≤0.8. These hits are highlighted in brown, MND1 is highlighted in blue. C) Gene ontology enrichment of screen hits from this study. D) Clonogenic outgrowth of HAP1 WT and three monoclonal ΔMND1 cell lines (blue) in response to increasing doses of IR. Data presented as mean±SEM, N=3. E) Clonogenic outgrowth of IR-treated HAP1 ΔMND1 cells overexpressing a doxycycline (Dox) inducible (GFP-)MND1 construct. Data presented as mean±SEM, N=3. F) Colony outgrowth of HAP1 cells treated with siRNAs targeting MND1 and HOP2 in response to increasing doses of IR, non-targeting (NT) siRNA as control. Data presented as mean±SEM, N=3. D)-F) Statistics depict p-values (* p ≤ 0.05, **** p ≤ 0.0001) after analysis using a Fisher’s test.

To narrow-down our hit list, we cross-referenced our screen with both replicates of a recently published IR screen in HAP1 cells (Francica et al., 2020). We applied the same cutoffs for all three IR datasets (p-value ≤0.05 and an odds ratio ≤0.8), which identified 37 common hits (Suppl. Figure 1A&B). This list includes genes known for their role in the response to IR like the Shieldin complex (Dev et al., 2018; Gupta et al., 2018; Noordermeer et al., 2018), *PRKDC* (DNAPKc; (Blackford and Jackson, 2017; Dong et al., 2018)) and *RNF168* (Kelliher et al., 2021), demonstrating that the screens were able to identify genes involved in the response to DNA damage. We also find several other genes including *CTDSPL2, PKM* and *RPRD2* with no previously described role in DNA repair. Interestingly, we identify *MND1* (highlighted in blue, Figure 1B), known as a meiotic recombination factor, but for which a general role in the DNA damage response in somatic cells has yet to be described.

MND1 is well known for its role during meiotic cross-over repair after DSB induction by the SPO11 nuclease (Gerton and Derisi, 2002; Henry et al., 2006). Interestingly, analysis of MND1 mRNA transcript levels shows expression in all analyzed tissue types to similar levels of RAD51 (Suppl. Figure 1C). This global expression across different tissue types is also observed for HOP2, the co-factor of MND1. Therefore, we conclude that MND1-HOP2 are ubiquitously expressed proteins. To address whether MND1 is involved in DNA repair in somatic cells, we first confirmed that loss of MND1 increases IR sensitivity in three independent HAP1 knockout clones (ΔMND1) (Figure 1D, Suppl. Figure 1D). Similarly, depletion of MND1 using either CRISPRi or siRNAs also causes increased sensitivity towards IR (Suppl. Figure 1E). We confirmed that the observed sensitivity towards IR in ΔMND1 cells is a direct result of loss of MND1, as exogenous expression of either MND1 or GFP-MND1 reduces IR sensitivity (Figure 1E, Suppl. Figure 1F). The partial rescue of IR sensitivity in this system correlates with a lower percentage of cells expressing the respective constructs (Suppl. Figure 1G). In summary, our haploid genetic screen identified MND1, which we here establish to have an important role in the response to IR in somatic cells.

In meiotic cells, MND1 is bound to a co-factor, HOP2 (Zhao and Sung, 2015; Zierhut et al., 2004). This MND1-HOP2 interaction is essential for the role of MND1 in the repair of SPO11-mediated DSBs during meiosis (Tsubouchi and Roeder, 2002). However, our screen did not identify HOP2 (gene name: *PSMC3IP*). Upon closer inspection, we found that gene-trap sense integrations in the *HOP2/PSMC3IP* locus were in fact decreased upon IR, but was only found significantly different when compared to 3 out of the 4 control datasets (Suppl. Table 1). When we depleted HOP2 using siRNAs, we indeed found a significant sensitization of HAP1 cells towards IR (Figure 1F). Furthermore, when we co-depleted MND1 together with HOP2, we found a similar sensitization as after MND1 depletion alone (Figure 1F). Similarly, siRNA-mediated depletion of HOP2 in HAP1 ΔMND1 cells also did not result in any increased sensitivity towards IR (Suppl. Figure 1H). This indicates that MND1 and HOP2 indeed act together in the response to IR in HAP1 cells. As MND1 and HOP2 were shown to interact together during meiosis (Tsubouchi et al., 2020; Zhao and Sung, 2015; Zierhut et al., 2004), we investigated whether they also interact in somatic cells. Co-immunoprecipitation (Co-IP) in cells expressing GFP-MND1 or GFP alone demonstrated that MND1 and HOP2 indeed interact in somatic cells in a DNA damage-independent manner (Suppl. Figure 1I). Together, these data confirm that MND1 and HOP2 act together in a complex during the response to IR in somatic cells, akin to their mutually dependent role during meiotic DNA damage repair.

After confirming that MND1 loss sensitizes HAP1 cells to IR, we aimed to confirm whether MND1 is important in the response to IR in different cell lines as well. The comparison of the LD50 of IR in control or MND1-depleted cells demonstrates that MND1 loss sensitizes U2OS, H1299, RPE1ΔP53 and, to a limited extent, HCT116, but not SAOS2 cells (Suppl. Figure 1J). MND1 and HOP2 are expressed in all cell lines tested (Suppl. Figure 1K). Hence, the lack of MND1-requirement in SAOS2 cells cannot be explained by differential expression of MND1 or HOP2. The only previously described role of MND1 in somatic cells is a role in alternative lengthening of telomeres (ALT) (Cho et al., 2014), a mechanism of telomere maintenance and extension that is closely related to HR (Pickett and Reddel, 2015; Sobinoff and Pickett, 2017). However, we do not find a correlation between the ALT status and IR-sensitization (HAP1, RPE1ΔP53 and HCT116 cells are ALT-; H1299, U2OS and SAOS2 cells are ALT+). Thus, ALT-status can also not explain the lack of MND1-requirement in SAOS2 cells. Collectively, we conclude that MND1 loss sensitizes most tested cell lines towards IR. Future work is necessary to understand the differential sensitivity of cell lines to loss of MND1 in response to IR-induced damage. This will shed more light on the dependencies of different genetic backgrounds to specific repair pathways.

### MND1 facilitates DSB repair by assisting in homologous recombination

After establishing a role for MND1 in the response to IR, we wanted to identify the specific mode of action of MND1. During meiosis, repair of SPO11-induced DSBs is critically dependent on the MND1-HOP2 complex, and the retention of DNA damage in the absence of MND1-HOP2 is well described (Gerton and Derisi, 2002; Petukhova et al., 2003; Zierhut et al., 2004). To test whether DNA DSBs are also retained in somatic cells, we tested whether MND1 is involved in the repair of IR-induced breaks in somatic cells. Hence, we first assessed DNA repair kinetics by quantification of 53BP1-foci as a proxy for the appearance and subsequent repair of DNA DSBs. For this, we used live-cell imaging of RPE1 cells in which we homozygously knocked-in a HALO-tag into the N-terminal site of the 53BP1-locus (Suppl. Figure 2A–E and see methods for cell line generation). Analysis of 53BP1 foci formation and resolution in asynchronously growing RPE1 cells revealed that MND1 depletion leads to slower repair and retention of DSBs after IR (Figure 2A, Suppl. Figure 2F&G), implicating MND1 in somatic DSB repair. However, when comparing the kinetics of 53BP1 foci resolution in MND1-depleted cells to BRCA1- or DNAPKc-depleted cells, we see a retention of DNA damage to a lesser extent. This indicates that MND1 has a less pronounced role in the DDR than BRCA1 or DNAPKc, which are known for their essential role in HR and NHEJ, respectively.

**Figure 2:**
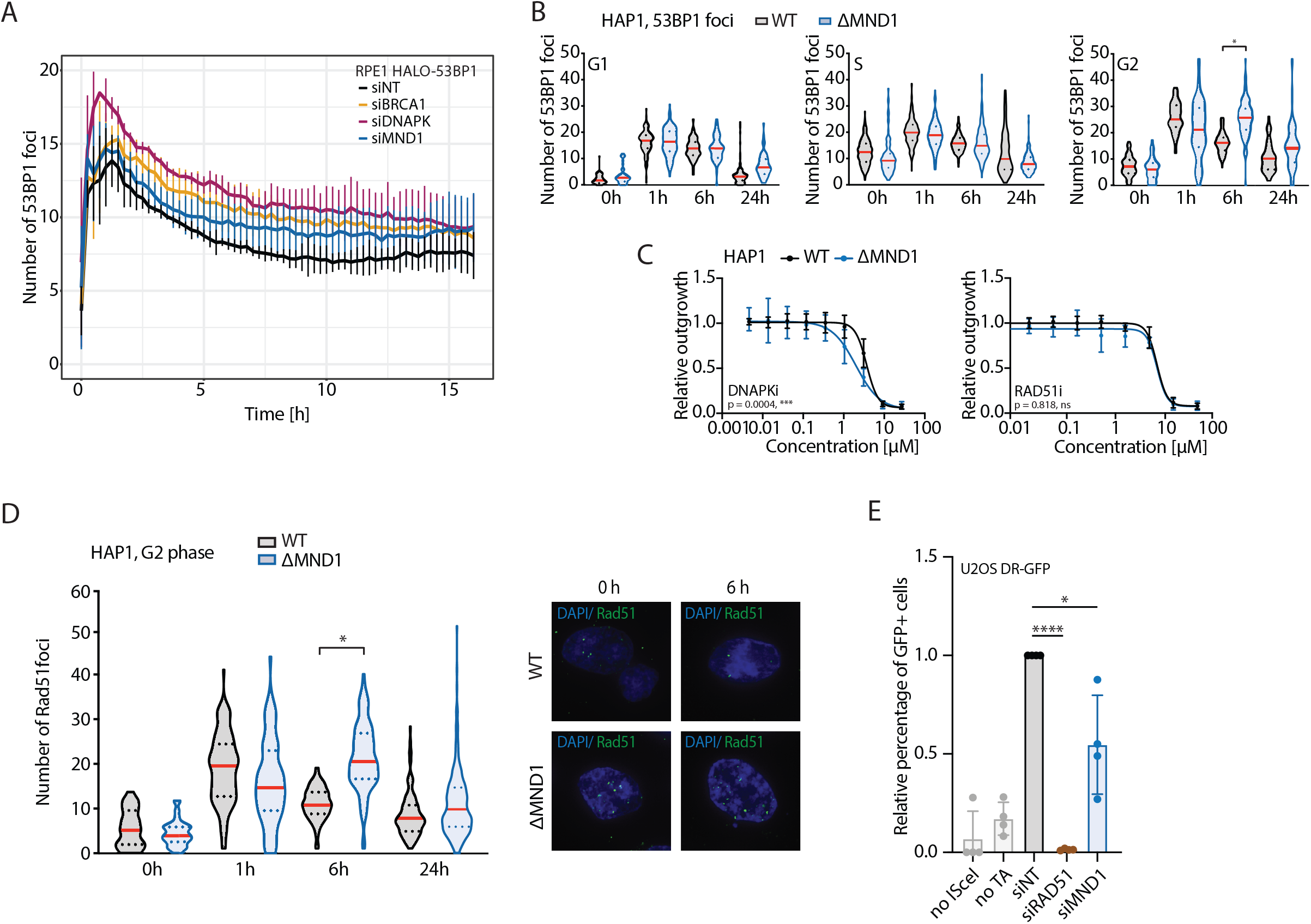
MND1 plays a role in the repair of DSBs via homologous recombination. A) 53BP1 foci formation in RPE1 HALO-53BP1_RPA1-mScarlet2 cells after treatment with 4Gy. Data presented as mean±SD, N=2. B) Immunofluorescence staining of 53BP1 foci in HAP1 WT and ΔMND1 cells during G1, S and G2 phases of the cell cycle at different time points after IR with 2Gy. Three independent replicates are displayed, statistical analysis (t-test) was performed on mean values (* p ≤ 0.05). C) Drug-response assays with DNAPKi NU7441 (N=4) and RAD51i B02 (N=3) in HAP1 WT and ΔMND1 cells. Data presented as mean±SD. Statistical analysis is performed on IC50 values of experiments (Suppl. Figure 2I). D) Immunofluorescence staining of RAD51 foci in HAP1 WT and ΔMND1 cells at different time points after 2Gy of IR. Three independent replicates are displayed, statistical analysis (t-test) was performed on the mean values of three experiments (* p ≤ 0.05). Example images are depicted on the right. E) HR frequency was assessed using the DR-GFP assay in U2OS cells treated with siNT, siRAD51 and siMND1. DSBs were induced by transfection of ISceI nuclease plasmid and triamcinolone acetonide (TA) addition. Data presented as mean±SD. Statistics depict p-values (* p ≤ 0.05, **** p ≤ 0.0001) after analysis using an unpaired t-test.

To explore if MND1 acts at different stages during the somatic cell cycle, we assessed 53BP1 focus formation and resolution in different cell cycle phases in HAP1 cells by immunofluorescence staining of 53BP1. Interestingly, loss of MND1 impaired 53BP1 foci resolution only in G2 phase (Figure 2B), suggesting that MND1 does not affect DNA repair in G1 and S phase. We confirmed that MND1 specifically affects DNA repair in G2 phase by analyzing yH2AX foci formation and resolution (Suppl. Figure 2H). Collectively, these data demonstrate that MND1 is involved in the repair of DSBs in the G2 phase of the mitotic cell cycle.

Following SPO11-induced break formation in meiosis I, the DSB is resected and MND1 acts to facilitate invasion of the single-stranded section of the DSB into the intact double-stranded homologous chromosome (Henry et al., 2006; Sansam and Pezza, 2015; Zickler and Kleckner, 2015). Given the established role of MND1 in meiotic recombination in germ cells, we next investigated if MND1 is involved in HR in somatic cells. First, we assessed the sensitivity of MND1 knockout cells to different DSB repair pathway inhibitors. We found that the loss of MND1 increases the sensitivity specifically towards DNA-PKcs inhibition, which inhibits DNA repair through NHEJ. Conversely, the sensitivity to inhibition of RAD51, and thus inhibition of HR, was unaltered in HAP1 ΔMND1 cells (Figure 2C, IC50 values of each replicate are depicted in Suppl. Figure 2I). The lack of sensitivity of ΔMND1 cells to RAD51 inhibitors implies that MND1 is redundant with the RAD51-dependent HR-repair. These data indicate that MND1 knockout cells rely on NHEJ, consistent with a (partial) deficiency in HR upon MND1 loss.

We next assessed at which step HR is compromised upon loss of MND1. To this end, we analyzed the appearance and disappearance of RAD51 foci after IR in the G2 phase. RAD51 is recruited to the DSB prior to strand invasion (Jasin and Rothstein, 2013), and MND1 facilitates strand invasion in meiotic cells (Bugreev et al., 2014; Petukhova et al., 2005). We find that RAD51 recruitment occurs with similar kinetics in ΔMND1 and WT cells (Figure 2D). However, the resolution of RAD51 foci occurs markedly slower in ΔMND1 cells (Figure 2D). This indicates that these cells exhibit a partial defect in the completion of HR, possibly at the level of strand invasion. To confirm that MND1 facilitates HR repair during the somatic cell cycle, we used the DR-GFP system to assess HR efficiency. In brief, induction of a DSB at a mutated, inactive, GFP can restore GFP fluorescence if that DSB undergoes HR-repair (Gunn and Stark, 2012). MND1 depletion caused a reduction of GFP+ cells, indicating that MND1 indeed facilitates HR in somatic cells (Figure 2E). Consistent with the notion that MND1 may facilitate some, but not all forms of HR-dependent repair, loss of MND1 led to a partial reduction in HR frequency, far less than the reduction obtained by depletion of RAD51.

HR in somatic cells is largely limited to S and G2 phase (Ceccaldi et al., 2016; Hustedt and Durocher, 2017), and therefore the reduction in HR observed after depletion of MND1 could be induced indirectly, through an altered cell cycle distribution in MND1-deficient cells. However, we show that the cell cycle distribution of U2OS cells depleted of MND1 is comparable to control cells (Suppl. Figure 2J), excluding cell cycle differences as a cause for the decrease in GFP+ cells. Thus, we conclude that MND1 plays an important and direct role in the repair of DSBs via HR in somatic cells.

### Loss of MND1 leads to increased sensitivity to some, but not all types of DSBs

Homologous recombination is an important repair mechanism at sites of endogenously induced DSBs. Ongoing replication is the source of the majority of endogenously occurring DNA lesions, most of which are repaired through HR. After establishing the role of MND1 in somatic HR after IR (Figure 2D), we aimed to assess whether this can be extended to other sources of DSBs, like replication-associated damage. We find significantly increased sensitivity towards the different DSB-inducers etoposide and doxorubicin (both topoisomerase II inhibitors), as well as the radiomimetic drugs neocarzinostatin (NCS) and zeocin (Figure 3A&B, Suppl. Figure 3B). By contrast, when we treated HAP1 ΔMND1 cells with inducers of replication stress, hydroxyurea (HU) and aphidicolin, there was little to no difference in the sensitivity of the MND1-deficient cells as compared to their wild-type counterparts (Figure 3A&B, Suppl. Figure 3A&B). Similar to replication stress induction, MND1 loss did not sensitize cells to camptothecin (CPT) (Figure 3A, Suppl. Figure 3B), a topoisomerase I inhibitor that creates DSBs specifically in S-phase, when a replication fork collides with the blocked topoisomerase I complex (Pommier et al., 2016). These data imply that MND1 is not required for the repair of DSBs that arise during DNA replication. To corroborate this finding we also tested if MND1 is involved in the repair of intra-strand DNA crosslinks (ICL), which often result in DSBs during replication (Deans and West, 2011; Semlow and Walter, 2021). We find no sensitization of ΔMND1 cells towards the ICL-inducing drugs cisplatin and mitomycin C (Figure 3A&B, Suppl. Figure 3A&B). This shows that MND1 is not involved in the HR-dependent repair of ICLs. At last, we treated cells with olaparib and talazoparib, two PARP inhibitors (PARPi) commonly used in the clinic. Comparing to data obtained in BRCA1/2-deficient cells, which are exquisitely sensitive to PARPi treatment (Noordermeer and van Attikum, 2019; O’Connor, 2015), we also observe a moderate sensitization towards olaparib and talazoparib after MND1 loss (Figure 3A&B, Suppl. Figure 3A&B). As PARPi treatment introduces PARP-trapping lesions that are converted to DSBs during S phase (Noordermeer and van Attikum, 2019), this indicates that there is some involvement of MND1 in the repair of replication-associated DSBs. However, we cannot exclude that this could be selectivity between DNA break structures having differential requirements for MND1. Taken together, these data show that the requirement of MND1 for the response of cells towards replication stress is limited to the highly HR-dependent PARPi-induced breaks.

**Figure 3:**
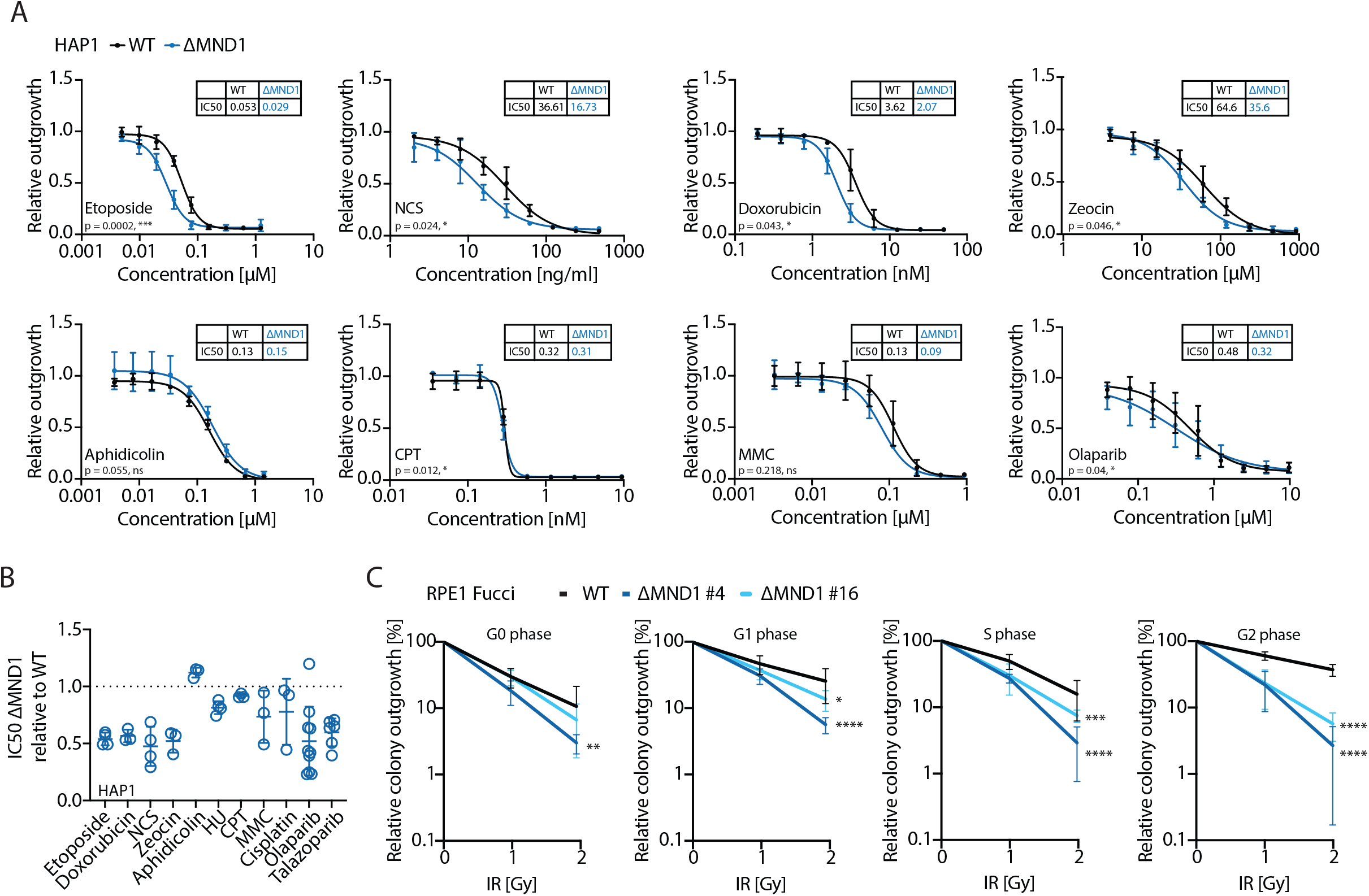
MND1 loss sensitizes to induction of a specific species of DSBs. A) Drug-response assays with etoposide (N=4), neocarzinostatin (NCS, N=4), doxorubicin (N=3), zeocin (N=3), aphidicolin (N=3), camptothecin (CPT, N=3), mitomycin C (MMC, N=3), olaparib (N=10) and talazoparib (N=6) in HAP1 WT and ΔMND1 cells. Data presented as mean±SD. Statistical analysis is performed on IC50 values (Suppl Figure 3B). B) Relative IC50 values of drug response curves from Fig2A and Suppl. Fig. 2A. IC50 values of ΔMND1 HAP1 cells are normalized to IC50 values of WT cells. Each dot represents a replicate experiment. C) Clonogenic outgrowth of RPE1 iCut Fucci WT and two ΔMND1 clonal cell lines after treatment with IR and sorting into different gates. Data presented as mean±SEM, N=6. Statistics depict p-values (* p ≤ 0.05, ** p ≤ 0.01, *** p ≤ 0.005, **** p ≤ 0.001) after analysis using a Fisher’s test.

We were intrigued by the differential requirement for MND1 in response to DNA damage that is or is not associated with replication. We reasoned that this difference could be because either I) the MND1-HOP2 complex is inhibited during S phase, and therefore not involved in the response to replication-associated breaks, II) differences in the repair of one- and two-ended DSBs that determine whether the MND1-HOP2 complex is necessary, or III) variability in the access of the repair template. To address the importance of MND1-HOP2 complex in various cell cycle stages after IR, we assessed the clonogenic outgrowth of RPE1 Fucci cells sorted from distinct cell cycle phases (cycling G1, S and G2 phase cells (Suppl. Figure 3C&D)). We observed significant sensitization in most ΔMND1 conditions compared to WT, with the exception of ΔMND1 clone 16 in G0 phase (Figure 3C). Strikingly, the further cells progressed through the cell cycle, the stronger the sensitization after MND1 loss became. Both G1 and S phase cells in this experiment showed significant sensitization after MND1 loss compared to WT. This indicates that either MND1 is involved in the response to a role at IR-induced DSBs inflicted during G1 and S phase, or that cells progress with DSBs from G1 and S phase further into G2, where MND1 is highly involved in the repair of the DNA damage. Notably, sensitization was most prominent in G2 phase (Figure 3C), consistent with our finding that depletion of MND1 affects repair in G2 cells only (Figure 2B). Interestingly, the lack of changes in DSB resolution we observed previously in G1 and S phases (Figure 2B) does not translate into a lack of sensitization of cells in these cell cycle phases (Figure 3C). Taken together, we assume that a carry-over of DSBs from G1 and S phase into G2 is the most likely explanation of our observation. Thus, our data show that the role of MND1 in DNA repair is most prominent in G2 phase cells and restricted to repair of two-ended DSBs.

### MND1 forms foci at DNA DSB sites together with yH2AX and RAD51

So far, we established that MND1 facilitates the HR repair of two-ended DNA DSBs during the mitotic cell cycle. However, we have not yet investigated whether MND1 localizes to DNA DSBs and is therefore directly involved in the repair of DSBs. For this, we visualized MND1 localization to sites of DNA damage by exogenous expression of a GFP-tagged MND1 fusion protein (GFP-MND1) in the RPE1 HALO-53BP1 cell line (Suppl. Figure 4A). We have previously demonstrated that this GFP-tagged MND1 construct rescues sensitivity in HAP1 cells to a similar extent as untagged MND1 (Figure 1E), warranting us to use this construct for further imaging experiments. GFP-MND1 readily forms foci after DNA damage induction as early as 2h after IR, and foci numbers continue to increase until they reach a plateau between 4h and 6h after DNA damage induction (Figure 4A). Since MND1 and HOP2 function epistatically (Figure 1E), we tested whether MND1 foci formation depends on HOP2. Interestingly, we see no foci formation after HOP2 depletion by siRNAs (Figure 4B and Suppl. Figure 4B), showing that MND1 foci formation is entirely dependent on its co-factor HOP2. These data show that MND1 is recruited to sites of DSBs, and that MND1 recruitment requires HOP2.

**Figure 4:**
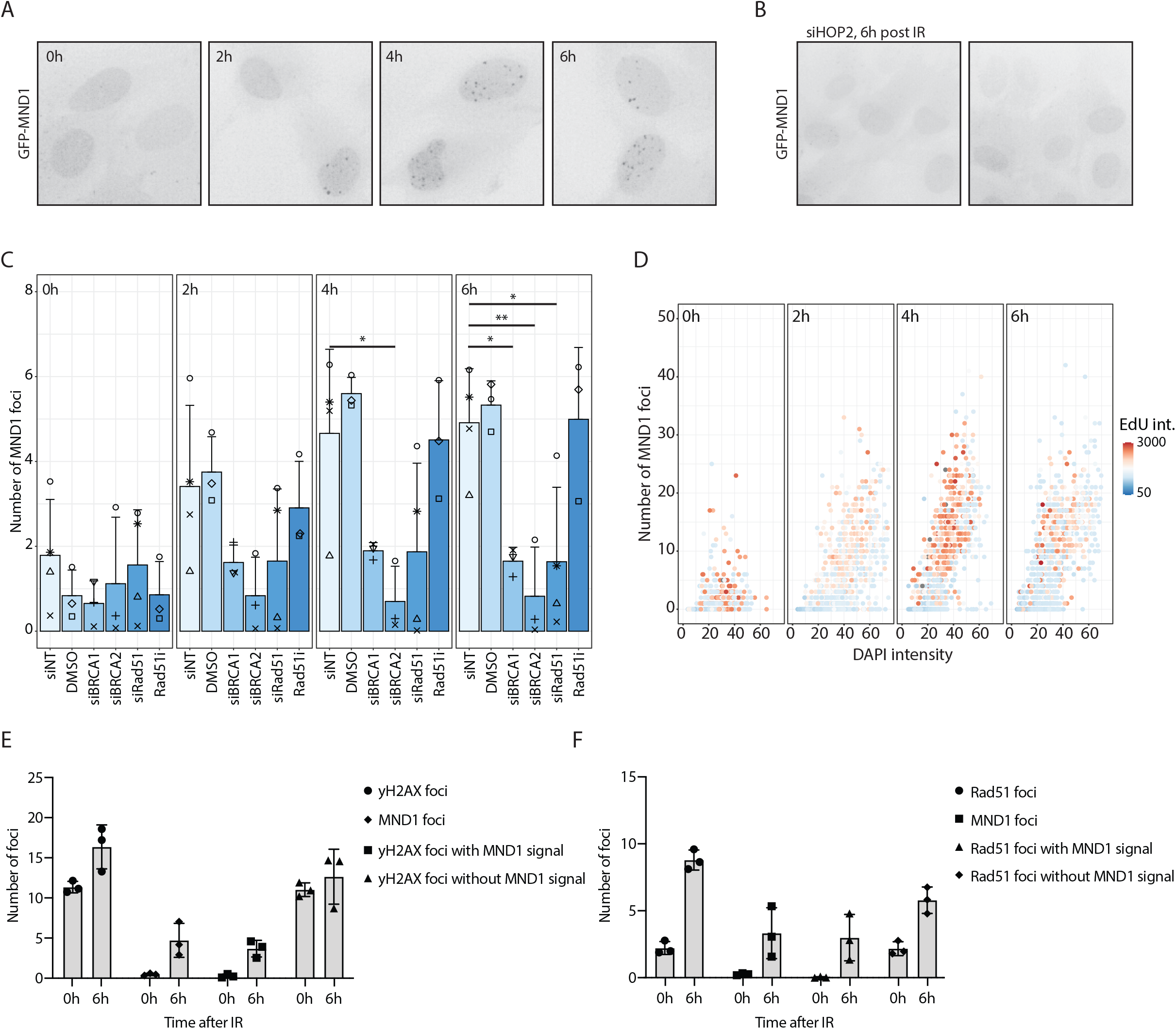
Localization of MND1 to a subset of yH2AX and RAD51-coated DSB locations is dependent on resection. A) Example images of RPE1 HALO-53BP1_RPA1-mScarlet2_GFP-MND1 cells after 4Gy of IR. Depicted are GFP-MND1 imaging examples at 0, 2, 4 and 6h after 4Gy. B) Example images of cells from A) treated with siHOP2 for 24h before IR with 4Gy. Images are taken 6h post IR. C) Number of GFP-MND1 foci detected at 0, 2, 4 and 6h after 4Gy IR. RPE1 HALO-53BP1_RPA1-mScarlet2_GFP-MND1 cells were treated with different siRNAs/inhibitors for HR pathway proteins. D) Co-staining of GFP-MND1 with DAPI and EdU to indicate the cell cycle phase-dependent formation of GFP-MND1 foci. This data is a representative experiment of two replicate experiments. E) Overlap of GFP-MND1 and yH2AX foci. Depicted are the number of either total yH2AX or MND1 foci, yH2AX foci with(out) MND1 signal, or MND1 foci with(out) yH2AX signal. F) Overlap of GFP-MND1 and RAD51 foci. Depicted are the number of either total RAD51 or MND1 foci, RAD51 foci with(out) MND1 signal, or MND1 foci with(out) RAD51 signal.

To further elucidate the notion that MND1 foci formation is dependent on resection of the DSB and ongoing HR repair, we depleted canonical HR factors using siRNAs. We find that MND1 recruitment is fully dependent on BRCA1, BRCA2 and RAD51 (Figure 4C). This places MND1 localization to sites of DSBs downstream of DNA end-resection and RAD51-loading, which is consistent with MND1’s role during meiosis (Chi et al., 2007). We also tested if the chemical inhibition of RAD51 would affect MND1 foci formation, using an inhibitor that specifically blocks sister chromatid exchange, but does not prevent RAD51 from binding to ssDNA (Huang and Mazin, 2014). Interestingly, we do still observe MND1 foci formation in this setting, which indicates that it is the presence of RAD51 at sites of ssDNA that is important for MND1 recruitment, whereas its functionality is dispensable. Therefore, we conclude that MND1 recruitment to sites of damage is dependent on the presence of RAD51-coated ssDNA, and occurs prior to RAD51-dependent invasion of the sister chromatid.

Consistent with the role of MND1 in the response to IR-induced DSBs in S and G2 phases of the cell cycle (Figure 3E), we find that MND1 localizes to DSBs in S and G2 phase (Figure 4D). Conversely, MND1 foci are not present in G1 phase. This indicates that MND1 is either involved in both S and G2 phase at sites of damage, or the loading of MND1 to DSBs during S phase is a preparation for when it is required later in G2.

Next, we aimed to analyze whether MND1 foci are formed at sites of DSBs, and more specifically, sites of HR repair. We find that virtually all GFP-MND1 foci are positive for yH2AX and RAD51 (Figure 4E&F and Suppl. Figure 4D), demonstrating that MND1 exclusively localizes to DNA DSB sites. However, only a fraction of IR-induced yH2AX- or RAD51 foci are coated with GFP-MND1, consistent with the fact that not all DSBs engage in HR and only a subset of HR breaks requires MND1 for their resolution (Figure 4E&F). This again indicates that, of the breaks that engage in RAD51-dependent HR, there is only a subset that depend on MND1 for repair.

Considering that MND1-HOP2 has a known role in HR at telomeres during ALT (Cho et al., 2014), we aimed to exclude that the GFP-MND1 foci we observe are only formed at telomeric DSBs at telomeres. For this, we used CRISPR/Cas9 to generate DSBs either at telomeres (‘Telo’) or in GAPDH pseudogenes (‘P63’, this crRNA generates around 18 DSBs in RPE1 cells in non-telomeric regions (Berg et al., 2019)). When we induced DSBs at these distinct locations, we could observe that even even though the overall induction of DSBs is higher when using the crRNA targeting telomeres (as shown by quantification of 53BP1 foci), the number of GFP-MND1 foci is smaller (Suppl. Figure 4E). This shows that the observed recruitment of GFP-MND1 to foci is not restricted to DSBs induced at telomeres, and strengthens our data showing a general role of MND1 in the HR repair of DSBs.

### MND1-deficient cells are prone to arrest at the G2 checkpoint after DSB induction

Thus far, we established that MND1 plays an important role in the repair of DSBs by aiding HR, and that loss of MND1 leads to increased DSB toxicity. It has been previously established that defects in DSB repair lead to stronger activation of cell cycle checkpoints via ATM and ATR, explaining growth arrest in those cells (Blackford and Jackson, 2017; Shaltiel et al., 2015; Shiloh, 2001). Specifically, it has been shown that HR defects lead to increased ATR-dependent cell cycle exit (Feringa et al., 2018). Thus, we were wondering whether the HR defects we observe after MND1 loss (Figure 2C–E) lead to increased cell cycle checkpoint activation. Indeed, we see a persistent increase in general DNA damage signaling in RPE1 ΔMND1 cells when assessing phosphorylation of H2AX (S139, yH2AX), CHK1 (S345) and CHK2 (T68), which demonstrates hyperactivation of ATM- and ATR-dependent checkpoint signaling upon loss of MND1 (Figure 5A). Consistently, we also observe a persistent increase in pCHK1-S345 in HAP1 ΔMND1 cells (Suppl. Figure 5A). This indicates that there are both unprocessed DSBs and resected ssDNA present (Blackford and Jackson, 2017), which shows that loss of MND1 is leading to increased checkpoint signaling which is potentially causing the growth arrest of ΔMND1 cells in response to DSB induction.

**Figure 5:**
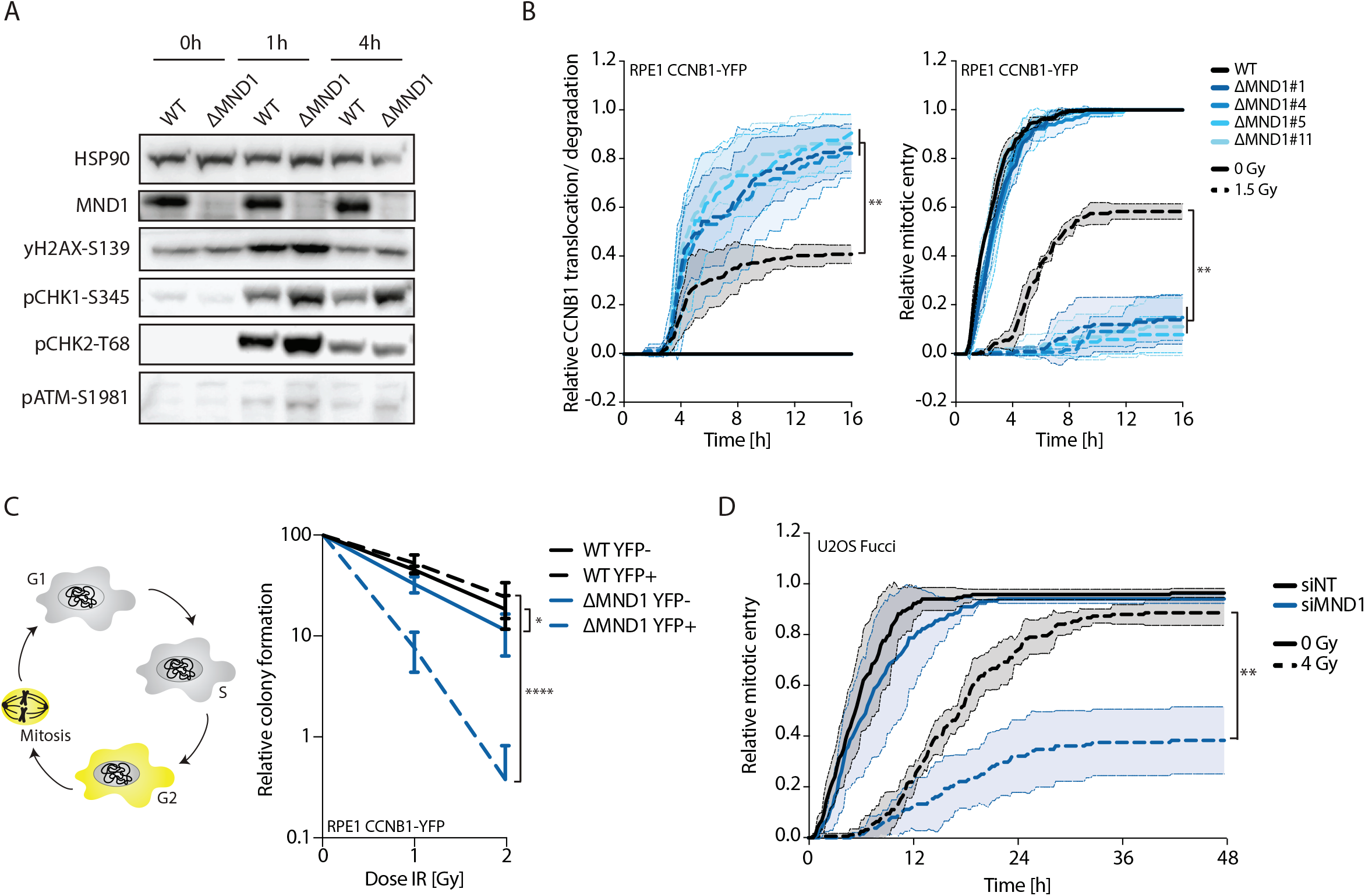
Loss of MND1 leads to hyperactivation of the G2 checkpoint and cell cycle arrest. A) Immunoblotting of checkpoint activation in RPE1 CCNB1-YFP WT and ΔMND1 cells. Blotting was performed at 0h, 1h or 4h after 4Gy of IR. B) RPE1 CCNB1-YFP cells were irradiated with 1.5Gy and CCNB1 translocation (left) or mitotic entry (right) is plotted over time in either WT or different ΔMND1 cell lines. Control cells that were not irradiated are depicted in full line. Data presented as mean±SD. Statistical analysis (t-test) is performed at 16h between WT and the separate ΔMND1 clones (** p ≤ 0.01). C) Colony outgrowth was assessed in RPE1 CCNB1-YFP WT and ΔMND1 cells after increasing doses of IR. Cells were sorted into YFP-(G1) and YFP+ (G2) cells after IR. Data is presented as mean±SD. Statistical analysis was performed using a Fisher’s test (* p ≤ 0.05, **** p ≤ 0.0001). D) Relative mitotic entry of G2 cells after 4Gy of IR. U2OS Fucci cells were treated with NT, MND1 and HOP2 targeting siRNAs and G2 cells were followed until mitotic entry. Data presented as mean±SD. Statistical analysis (t-test) is performed at 48h (** p ≤ 0.01, *** p ≤ 0.001, **** p ≤ 0.0001).

We have previously shown that HR intermediates can trigger a permanent cell cycle exit in G2 phase (Feringa et al., 2018). This is because unresolved RPA and RAD51-coated DSBs elicit a strong G2 checkpoint response which leads to nuclear translocation or degradation of cyclin B1, a marker of irreversible cell cycle arrest (Charrier-Savournin et al., 2003; Krenning et al., 2014). We reasoned that HR intermediates that are left unresolved in ΔMND1 cells are causing a strong arrest specifically in G2 phase to prevent cells from entering mitosis. To test is MND1 loss induces a G2 arrest, we made use of an RPE1 cell line with endogenously tagged cyclin B1 (RPE1 CCNB1-YFP; (Krenning et al., 2014; Shaltiel et al., 2014)), where we knocked-out MND1 (Suppl. Figure 5B). Indeed, cells deficient for MND1 show a dramatic increase in nuclear translocation or degradation of cyclin B1, and a corresponding decrease in mitotic entry (Figure 5B). This is also evident when we deplete MND1, and its co-factor HOP2, by siRNA treatment (Suppl. Figure 5C). This demonstrates that the loss of MND1 potentiates growth arrest by IR in G2 phase.

As we have now seen MND1 loss to infer a dramatic reduction in mitotic entry, co-occurring with a G2 checkpoint activation, we tested whether G2 cells in this system are more sensitive towards IR than G1 cells. For this, we irradiated WT and ΔMND1 RPE CCNB1-YFP cells and then sorted both a YFP- and YFP+ population for clonogenic outgrowth (G1 and G2 phase cells respectively, Suppl. Figure 5D). We indeed find that the G2 (YFP+) ΔMND1 cells are specifically sensitive towards IR when compared to WT cells (Figure 5C). These data are consistent with a role for MND1 in HR during G2 phase and its loss resulting in defects in strand invasion, producing persistent HR intermediates that can drive a permanent cell cycle arrest after DNA damage.

Lastly, we tested if the loss of MND1 also limited the proliferation of transformed cells that receive DSBs in G2. For this, we imaged U2OS Fucci cells that were depleted of MND1 using siRNAs. When we followed single G2 phase cells entering mitosis, we confirmed that loss of MND1 reduces mitotic entry specifically upon IR (Figure 5D). We conclude that loss of MND1 results in a stronger G2 checkpoint, thereby increasing radiation sensitivity.

## Discussion

Here we identified for the first time a previously unrecognized role for the MND1-HOP2 complex during somatic HR. We show that cells lacking MND1 demonstrate increased sensitivity towards DNA damage, specifically y-irradiation and drug-induced two-ended DSBs. Similar to meiosis, where MND1 and HOP2 bind RAD51 and facilitate strand invasion and D-loop formation (Bugreev et al., 2014; Petukhova et al., 2005), we demonstrate the importance of MND1 for efficient HR in somatic cells. Furthermore, complex formation of MND1-HOP2 is necessary for the localization of MND1 to sites of (RAD51-covered) DSBs, where it facilitates repair.

When we depleted MND1 in multiple cell lines, we found a variable effect of sensitization (Suppl. Figure 1J). We have yet to identify the underlying reason for differences between cell lines. One hypothesis is that the MND1-HOP2 complex is specifically active in G2 phase cells, where we also see the strongest sensitization in sorted RPE1 cells. As MND1 was not previously identified having a general role in somatic cells, we hypothesize that previous studies performed in asynchronous cells have masked this cell cycle phase specific role of MND1. This leads us to speculate that there are still other genes with unidentified roles in the DDR with such a specific involvement in DNA repair.

The formation of GFP-MND1 foci upon IR (Figure 4) establishes the direct involvement of MND1 at sites of damage. We confirmed that the recruitment of MND1 to DSBs is entirely dependent on HOP2 and the initiation of HR, up to successful RAD51 loading (Figure 4B&C). This is in line with previous experiments performed in vitro, where MND1 binds to established RAD51-coated ssDNA, to thereby aid in D-loop formation (Bugreev et al., 2014; Chi et al., 2007; Petukhova et al., 2005). The observed localization of GFP-MND1 to sites of DSBs is in contrast to previous studies, where in yeast meiotic cells MND1 foci were found to be formed randomly throughout the nucleus, rather than at sites of RAD51 foci (Tsubouchi and Roeder, 2002; Zierhut et al., 2004). The difference between our observation and previously published data could be underlying differences in MND1 usage between yeast and mammalian cells as MND1 and HOP2 are only in mammalian cells expressed during both the meiotic and the mitotic cell cycle. Therefore, we propose that there are critical differences in the function of MND1 in its localization to sites of DSBs between yeast and mammalian systems, and it would be interesting to study whether MND1 is recruited to RAD51-covered DSBs in mammalian meiotic cells.

We show that during somatic DSB repair, MND1 is readily recruited to DSBs covered with both yH2AX and RAD51 (Figure 4E&F). However, not all yH2AX or RAD51 foci recruit MND1 (Figure 4E&F). This goes together with our data showing that I) MND1 loss is less detrimental to 53BP1 foci resolution than depletion of BRCA1 (Figure 2A) and II) the HR-efficiency in the DR-GFP assay is reduced to ~50%, whereas RAD51 depletion leads to full loss of HR. This indicates that MND1 is required for repair of only a subset of DSBs engaged in HR. However, it is possible that we do not observe full overlap of RAD51 foci with MND1 as these experiments are not stratified for cell cycle. We hypothesize that MND1 foci in S phase localize less to RAD51 than in G2 phase, because even though MND1 can localize to foci in S phase, its role in repair during S phase is minor (foci resolution is unchanged and S phase cells are less radiosensitive than G2 cells, Figures 2B and 3C). Further experiments have to be conducted to establish the role of MND1 in S phase compared to G2 phase and to identify a determining factor for the conditions and types of breaks that require the MND1-HOP2 complex for their repair.

We have uncovered that not all HR-mediated repair requires MND1, which leads to speculate what the differences of these DSBs are. Based on our observations, we hypothesize that MND1 is required for repair of only two-ended DSBs, and not one-ended DSBs. Strikingly, the largest sensitization to IR upon MND1 loss was observed when DSBs were induced in G2, outside the context of replication. Moreover, when DSBs were generated by replication stress inducers aphidicolin, CPT or HU, only a slightly increased sensitization could be observed after HU treatment only (Figure 3A&B). During meiotic recombination, the homologous chromosomes used for crossovers are already compacted and apart from each other, calling for the requirement of efficient search of the homologous region. This has been reported to require the presence of the MND1-HOP2 complex (Pezza et al., 2010). We propose that our observed S phase sensitization to IR is carry-over of DSBs from G1/S phase into G2, where the role of MND1 is the most prominent. We speculate that this reflects a specific role of the MND1-HOP2 complex in the repair of DNA damage, where replication has concluded and the DNA strands are closed and sister chromatids taken apart, similar to the role of the MND1-HOP2 complex in meiosis. Therefore, more complex methods for homology search are necessary, spanning longer distances in the nucleus. We envision that DSBs after successful replication without the sister chromatid in direct proximity require an advanced mechanism for homology search and strand invasion. The lack of sensitization of ΔMND1 cells towards classical replication stress inducers supports this model. Furthermore, both MND1 and HOP2 are non-essential in human somatic cells. This is in striking contrast to other, ‘classical’, HR factors like RAD51, BRCA1 and BRCA2, which are involved in the repair of replication-associated DSBs as well (Bonilla et al., 2020; Takaoka and Miki, 2018; Wassing and Esashi, 2021), are essential factors for cell survival.

What makes this specific role of the MND1-HOP2 complex so interesting, is that these proteins are by themselves dispensable for normal cell survival. As MND1-HOP2 are not involved in the replication-associated repair of DSBs, and consequently not involved in the repair of most endogenous DNA damage, their loss is dispensable for cellular survival under normal growth conditions. However, their loss renders cells highly sensitive to induction of exogenous damage by e.g., IR or TOP2i (as seen in Figures 1 and 3). Therefore, the interference with MND1-HOP2 function in cells could be a potentially interesting approach for cancer combination treatment with targeted DSB induction. This has the potential to be an efficient treatment, specifically in G2-phase rich tumors.

## Acknowledgments

We thank members of the Medema lab for helpful discussions and input on the manuscript. We want to further thank the flow cytometry and digital microscopy facilities for technical support. This work was supported by funding from the Oncode Institute. The Oncode Institute is partly supported by KWF Dutch Cancer Society.

## Author Contributions

L.Ko., L.Kr. and R.H.M. conceived and designed the study. L.Ko., A.F., L.v.B., E.S.K. and M.S. performed experiments, data processing and figure preparation. B.v.B. wrote the Fiji plugin for image processing and analysis. The screen was performed and analyzed by F.M.F. and V.A.B. under the supervision of T.R.B. L.Ko., L.Kr. and R.H.M. wrote the manuscript with input from all authors.

## Declaration of Interests

The authors declare no competing interest.

## Material and Methods

### Cell culture

Human-derived derived near-haploid HAP1 cells were cultured in IMDM (GIBCO) supplemented with 12% FCS, 1% GlutaMAX (GIBCO)) and 100 U/ml penicillin-streptomycin. RPE1 (CCNB1-YFP, Fucci and ΔP53), HCT116 and SAOS2 cells were cultured in DMEM:F12 (GIBCO) supplemented with 6% FCS, 1% GlutaMAX (GIBCO) and 100 U/ml penicillin-streptomycin. U2OS DR-GFP (Gunn and Stark, 2012) and Fucci cells were cultured in DMEM (GIBCO) supplemented with 6% FCS, 1% GlutaMAX (GIBCO) and 100 U/ml penicillin-streptomycin. H1299 cells were cultured in RPMI 1640 (GIBCO) supplemented with 12% FCS, 1% GlutaMAX (GIBCO), 1% HEPES (GIBCO), 1% MEM non-essential amino acids (GIBCO) and 100 U/ml penicillin-streptomycin.

### Haploid genetic screen

Genes essential for the fitness of cells treated with y-irradiation were identified as previously described (Blomen et al., 2015). In brief, gene trap retrovirus was produced in HEK293T cells. After harvesting the virus, approximately 40 million HAP1 cells were mutagenized. The mutagenized cells were treated with y-irradiation (1Gy, every other day) and passaged for 10 days in total. After passaging, cells were collected and fixed. Fixed cells were stained with DAPI to allow sorting for haploid cells only. The genomic DNA was isolated using a DNA mini kit (QIAGEN). The gene trap insertion sites were amplified by LAM-PCR and sequenced using primers containing Illumina adapters (Blomen et al., 2015). Mapping and analysis of insertions sites is described in detail by Raaijmakers et al. (Raaijmakers et al, 2018). In short, sequence reads were aligned to the human genome (hg19) to obtain the genomic locations of insertion sites. Subsequently, the gene-trap insertions were intersected with Refseq gene coordinates to ascertain intragenic integrations and the orientation with respect to the transcriptional direction of the gene. Overlapping gene regions that introduce ambiguity to insertion site direction calling were disregarded. To identify genes that are affecting fitness in IR-treated cells, the sense and antisense orientation integrations for each were compared to those in four independent published untreated datasets ((Blomen et al., 2015); NCBI SRA accession no. SRP058962) using a Fisher’s exact test.

### Ionizing radiation and clonogenic outgrowth

Cells were irradiated using a Gammacell Exactor (Best Theratronics) with a ^137^Cs source. For assessing the sensitivity of cell lines towards y-irradiation, low amounts of cells were plated per well, treated with different doses of irradiation and grown into single colonies for six days. The number of colonies was then counted and normalized to the untreated condition.

### Go term enrichment analysis

Hits from the haploid genetic screen were analyzed for gene ontology (GO) enrichment using http://geneontology.org/ (release 2022-07-01: 43.558) (Carbon et al., 2021; Gene and Consortium, 2000).

### siRNA and crRNA transfections

siRNA transfections were performed using RNAiMAX (Invitrogen) according to the manufacturer’s guidelines. The following siRNAs were used in this study: siNT (Non-targeting; Dharmacon), siMND1 (Dharmacon), siPSMC3IP (HOP2; Dharmacon), siRAD51 (Dharmacon), siBRCA1 (Dharmacon), siBRCA2 (Dharmacon), siPRKDC (DNAPKc; Dharmacon) and siTP53 (p53; Dharmacon).

A crRNA targeting exon 4 of MND1 (ΔMND1 cell generation, 5’-CAAGTAAAGCTCTTCATGCA-3’), GAPDH pseudogenes (P63, 5’-AACGGGAAGCTTGTCATCAA-3’) (Berg et al., 2019) or telomeres (Telo, 5’-UUAGGGUUAGGGUUAGGGUU-3’) (McCaffrey et al., 2017) were transfected using RNAiMAX (Invitrogen). In brief, 20nM crRNA and tracrRNA are incubated with RNAiMAX (Invitrogen) in OptiMEM (Gibco). After 20min of incubation, the transfection mix is added to the cells. Indel generation in the MND1 locus is validated by TIDE (Brinkman et al., 2014) analysis of PCR products.

### CRISPRi mediated knockdown

Two MND1 sgRNAs (#1 5’-GCGGCGAAGCCCACACACTA-3’, #2 5’-GCTGCGCCCGCGCCATGGTA-3’) targeting the promoter were cloned into a pLV-sgRNA plasmid and lentivirally transduced into cells.

### Clonogenic outgrowth assay/ drug response assays

To assess colony outgrowth after irradiation, 250 cell were seeded per well in 6-well plates. Cells were fixed after 7 days of growth in 80% methanol and stained with 0.2% crystal violet. Colonies were counted and normalized to the unirradiated control.

For drug response assays, 500 cells were plated per well in 96-well plates. After treatment with various drugs, cells were grown for 7 days. Cells were fixed in 80% methanol and stained with 0.2% crystal violet. Crystal violet staining was analyzed after treatment with 10% acetic acid in water. Intensity of staining was then quantified using a Biotek Epoch Microplate Reader.

### Endogenously-tagging of 53BP1 and RPA1

CRISPR/Cas9 was used to create endogenously-tagged RPE1 iCut cell lines with HALO-53BP1 and RPA1-mScarlet2. For tagging endogenous 53BP1 N-terminally with HALO, a 3x-HA-HALO-tag flanked with 200bp homology arms was acquired in a gBlock. RPA1 was tagged at the C-terminal site with the mScarlet2 fluorophore, a mScarlet2-3x-HA flanked with 90bp homology arms was acquired in a gBlock. A mutation in the PAM sequence was introduced to prevent re-cutting by pSpCas9 after initial break repair. The gBlocks were cloned into a KpnI-XbaI digested pUC19 vector (Addgene, #50005) by Gibson assembly. To insert the tags, CRISPR/Cas9-induced DSBs were generated using the pSpCas9(BB)-2A-Puro (PX459) plasmid (addgene #62988) expressing the following sgRNAs that were designed in close proximity to or overlapping with the start codon of 53BP1 and the stop codon of RPA1 using CRISPOR and BLAST: 53BP1 5’-GAGCGCGAGGGACCTCCCGCC-3’ and RPA1 5’-GAGAAGTGCATTGATGTGAG-3’. For nucleofection, RPE-1 iCut cells were plated and treated for 48h with p53 siRNA (see siRNA transfections), M3814 (MCE, 1mM), doxycycline (Sigma, 1mM) and SHIELD-1 (Aobious, 1μM). Nucleofection was performed using a Amaxa Nucleofector treating cells with 1ug pUC19 plasmid, 1 ug PX459 plasmid and blasticidin-expressing plasmid in Amaxa buffer (Lonza) with 100 mM ATP. After nucleofection, cells were plated and selected with blasticidin (10μg/ml) for 48h. To generate monoclonal endogenously tagged cell lines, successfully nucleofected cells were identified by FACS (BD FACSAria). HALO-53BP1 cells were incubated with a HALO-ligand, following the ligand labelling protocol. RPA1-mScarlet2 cells were identified by red fluorescence. Single clones were grown out and tested for endogenous knock-in by performing PCRs outside homology arms: OH-ARM_53BP1 FWD 5’-TCCATGCTGCCATGGAAACG-3’ and REV 5’-AATCTGTTCGCCAGAGGCCC-3’, HALO-FWD 5’-ATGGCAGAAATCGGTACTGG-3’

### Immunofluorescence staining and fixed-cell imaging

Cells were pre-extracted using 0.5% Triton X-100 in PBS on ice for 30s and immediately fixed on coverslips for 15 min at room temperature (RT) using a final concentration of 3.5% formaldehyde. Then cells were permeabilized for 5 min using 0.5% Triton X-100 in PBS. Cells were blocked in PBS supplemented with 0.1% Tween-20 (PBS-T) with 5% bovine serum albumin (BSA) for 1h. Primary antibody incubation was performed at RT for 1.5h (antibodies and dilutions stated below). Coverslips are washed with BSA in PBS-T and secondary antibody incubation is performed at RT for 1h. After incubation with secondary antibody, coverslips were washed with PBS. EdU was stained by incubation in EdU staining buffer (100 mM Tris-HCl pH8.5, 1mM CuSO4), with 100 mM ascorbic acid and AF-647 azide (Invitrogen, 1/1000) for 30 min at RT. After washing 3 times with PBS-T coverslips were mounted on microscopic slides using Prolong Gold (Invitrogen) and stored at 4°C.

### Immunofluorescence and live-cell imaging

Cells were either fixed and stained as described above or grown in Lab-Tek II chambered coverglass (Thermo Scientific) in tissue culture medium outfitted with a CO2 controller set at 5%. Images for Figure 2 were obtained using a DeltaVision Elite (Applied Precision) maintained at 37°C and 5% CO2 equipped with a 40x and 63x PLANApo S lens (Olympus) and cooled CoolSnap CCD camera. Images for Figure 4 were obtained using a THUNDER Imager (Leica Microsystems) maintained at 37°C and 5% CO2 equipped with a 63x/1.40-0.60 OIL Obj. HC PL APO objective and a deep-cooled 4.2 MP sCMOS camera.

### Foci quantification

For foci quantification in Figure 2, a previously published ImageJ macro was used (Feringa et al., 2016).

For foci quantification in Figure 4, images were split into single timepoints. Nuclear foci were quantified in Fiji (Schindelin et al., 2012), using a custom-built ImageJ macro that enabled automatic and objective foci analysis (https://github.com/BioImaging-NKI/Foci-analyzer). Initially, cell nuclei were detected by thresholding the (median-filtered) DAPI signal, followed by a watershed operation to separate touching nuclei. In a later version StarDist (Schmidt et al., 2018) was used for nuclei segmentation.

Brief outline of the foci detection workflow (in 2D): After maximum intensity z-projection, the foci signal is background-subtracted using a Difference-of-Gaussians filter. Foci candidates are identified as local maxima exceeding a user-adjustable threshold. These maxima are then used as seeds for MorpholibJ’s (Legland et al., 2016) marker-controlled watershed segmentiation, executed on the GPU using CLIJ2/CLIJx (Haase et al., 2020), followed by size filtering to exclude very small foci. Overlay images of segmented nuclei, detected foci and original signals provide a convenient way to inspect the results and optimize parameters, depending on foci size, intensity and noise levels. In experiments with two foci channels, foci are considered co-localized if their spatial overlap is at least 1 pixel.

### Western Blot

Western Blot analysis was performed as previously described (Feringa et al., 2016). In brief, proteins were separated using SDS-polyacrylamide gel electrophoresis and transferred to nitrocellulose membranes. Membranes were blocked in 5% BSA in PBS-T and afterward incubated with primary antibodies overnight at 4°C (dilutions of antibodies indicated below). Secondary antibodies are incubated as stated below for 1h at room temperature. Proteins were visualized using enhanced chemiluminescence (ECL) (GE Healthcare).

### Antibodies and chemicals

The following primary antibodies were used in this study: anti-H2AX ser139 (yH2AX, 05–636 Upstate, 1:500 IF, 1:1000 WB), anti-RAD51 (ab63801, abcam, 1:500 IF), anti-pCHK1 ser345 (2344, Cell Signaling, 1:500 WB), anti-pCHK2 thr68 (2661, Cell Signaling, 1:1000 WB), anti-GFP (11 814 460 001, Roche, 1:1000 WB/IF), anti-HA (H9658, Sigma, 1:1000 WB), anti-histone H3 (ab1791, abcam, 1:1000 WB), anti-HSP90 (sc7947, Santa Cruz, 1:1000 WB), anti-MND1 (235395, abcam, 1:1000 WB), anti-HOP2 (11339-1-ap, Thermo, 1:1000 WB), anti-pATM ser1981 (#4526, Cell Signaling, 1:1000 WB).

The following secondary antibodies were used for western blot experiments: peroxidase-conjugated goat anti-rabbit (P448, DAKO, 1:1000) and goat anti-mouse (P0447, DAKO, 1:1000). Secondary antibodies used for immunofluorescence were goat anti-mouse-Alexa 488 (A11029, Mol probes, 1:1000) and goat anti-rabbit-Alexa 568 (A11011, Mol probes, 1:1000). Chemicals used in this study: DNAPK inhibitor (NU-7441; 14881, Cayman), olaparib (AZD2281, M1664, bioconnect), talazoparib (AZD2461, SML1858, Sigma), RAD51 inhibitor (B02, 553525, Merck), neocarzinostatin (NCS, N9162, Sigma), doxorubicin (D1515, Sigma), etoposide (E1383, Sigma), zeocin (R250-01, Invitrogen), mitomycin c (MMC, M0503, Sigma), cisplatin (P4394, Sigma), hydroxyurea (HU, 0210202310, MP biomedicals), aphidicolin (A0781, Sigma), camptothecin (CPT, 0215973225, MP biomedicals).

### Flow cytometry analysis and sorting

Cells were trypsinized and resuspended in PBS supplemented with 1% FCS for sorting, using a BD FACSAria Fusion. G2 cells were sorted based on Cyclin B1-YFP signal and replated for colony counting or harvested for propidium iodide (PI) staining. RPE1 Fucci cells were sorted based on Azami-Green and Kusabira-Orange signal depending on cell cycle phase (gating strategy indicated in Suppl. Figure 3C), and replated for colony counting.

GFP+ and BFP+ cells were analyzed using BD LSRFortessa after cells were harvested in trypsinization and resuspended in PBS + 1% FCS. Cell cycle distribution was analyzed by staining fixed cells with PI.

### DR-GFP assay

U2OS DR-GFP cells were gifted by the Stark lab and the assay was performed as described in (Gunn and Stark, 2012). In brief, 72h after siRNA transfection (see above), media was changed to antibiotics-free DMEM media, and U2OS DR-GFP cells were then transfected with 1μg I-SceI-RFP expressing plasmid using Lipofectamine 2000. 3h post transfection, media was changed and supplemented with triamcinolone acetonide (TA) to induce nuclear translocation of the ISceI protein. After 72h, cells were harvested and RFP and GFP positivity were measured using a BD LSRFortessa.

---

**Suppl. figure 1:**
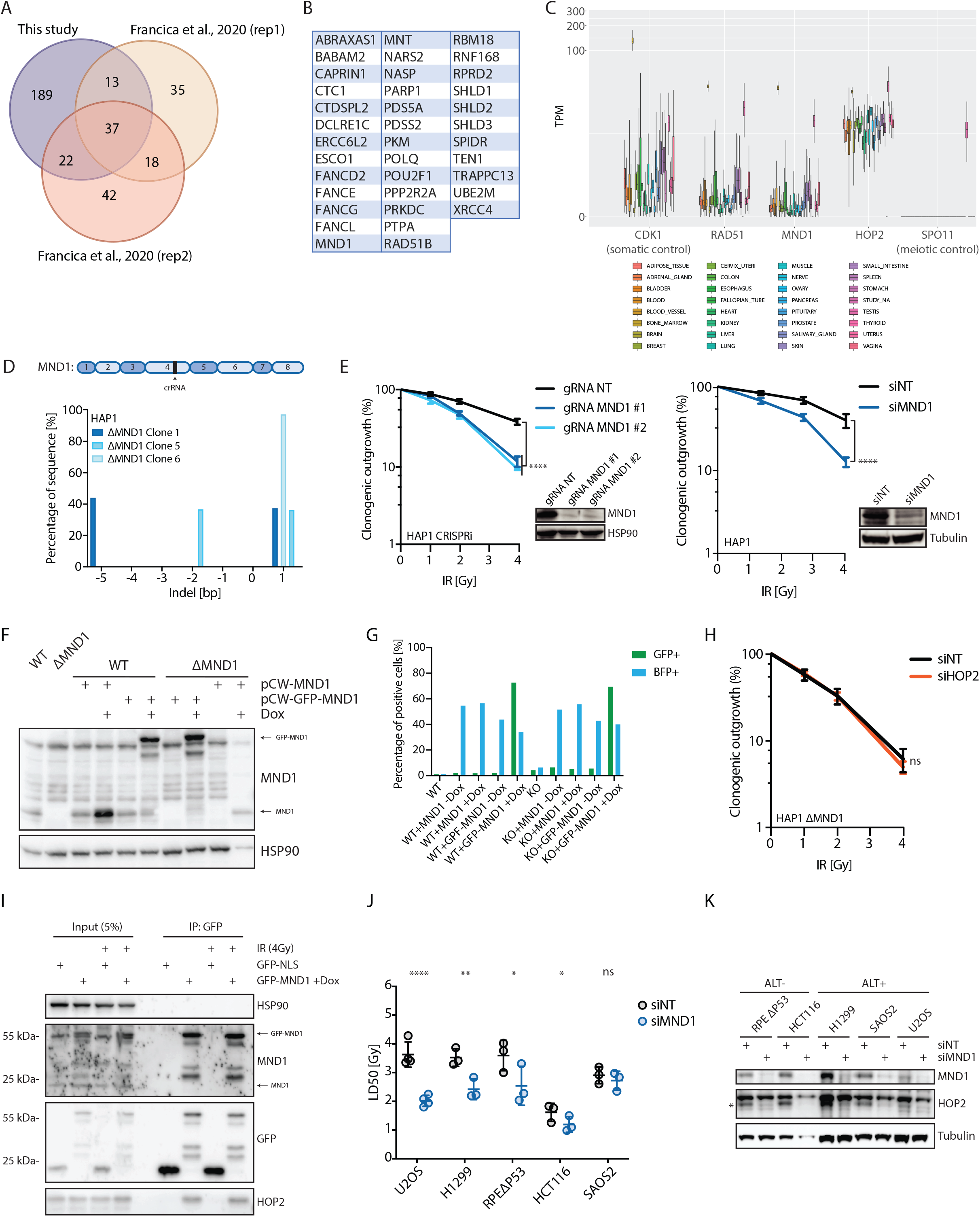
Confirmation of IR sensitivity phenotype. A) Screen hits of this study were selected to have an FDR-corrected p-value ≤ 0.05 when comparing the IR screen to four independent WT datasets, and an odd-ratio ≤ 0.8. The same criteria were used to filter hits from a previous study (Francicia et al., 2020) with a similar treatment regimen, performed in HAP1 cells as well. Depicted is an overlap of hits from all three replicates of screens. B) List of 10 overlapping hits found in all three screens (left). On the right is a list of 13 hits found overlapping in two of the three screens C) MND1 and HOP2 expression across tissues in GTEx data. D) MND1 exon structure, indicating the crRNA targeting site in Exon 4. Below is depicted the Indel pattern of three monoclonal ΔMND1 cell lines. E) Colony outgrowth of HAP1 CRISPRi cells expressing two different sgRNAs against MND1 promotor (left) and HAP1 cells treated with NT and MND1 siRNAs (right) in response to increasing doses of IR. Immunoblotting of MND1 and Tubulin protein is depicted on the side. Data presented as mean±SEM, N=3. Statistical analysis was performed using a Fisher’s test (**** p ≤ 0.0001). F) Immunoblotting for MND1 and HSP90 in HAP1 WT and ΔMND1 expressing pCW_(GFP-)MND1 constructs ±Doxycycline addition. G) Flow cytometry analysis of HAP1 WT and ΔMND1 expressing (GFP-)MND1 constructs ±Doxycycline addition to assess GFP and BFP expression. H) Colony outgrowth of HAP1 ΔMND1 cells treated with NT and HOP2 targeting siRNAs in response to increasing doses of IR. Data presented as mean±SEM, N=3. I) Immunoprecipitation (IP) experiment in HAP1 WT cells expressing either a GFP-NLS or a GFP-MND1 construct. IP was performed after 4Gy of IR, with beads recognizing GFP. Immunoblotting shows a pull-down of GFP-MND1 (with MND1 and GFP antibody) and HOP2 binding. J) Clonogenic outgrowth data of U2OS, H1299, RPEΔP53, HCT116 and SAOS2 cells after treatment with siNT or siMND1 in response to IR. Depicted are IC50 values of the dose of IR presented as mean±SEM, N=3. K) Immunoblotting of MND1 and HSP90 protein in different cell lines (RPE ΔP53, HCT116, H1299, SAOS2, U2OS) after siRNA-mediated depletion of MND1. A)-C)&J) Statistics depict p-values (* p ≤ 0.05, ** p ≤ 0.01, **** p ≤ 0.0001) after analysis using a Fisher’s test.

**Suppl. figure 2:**
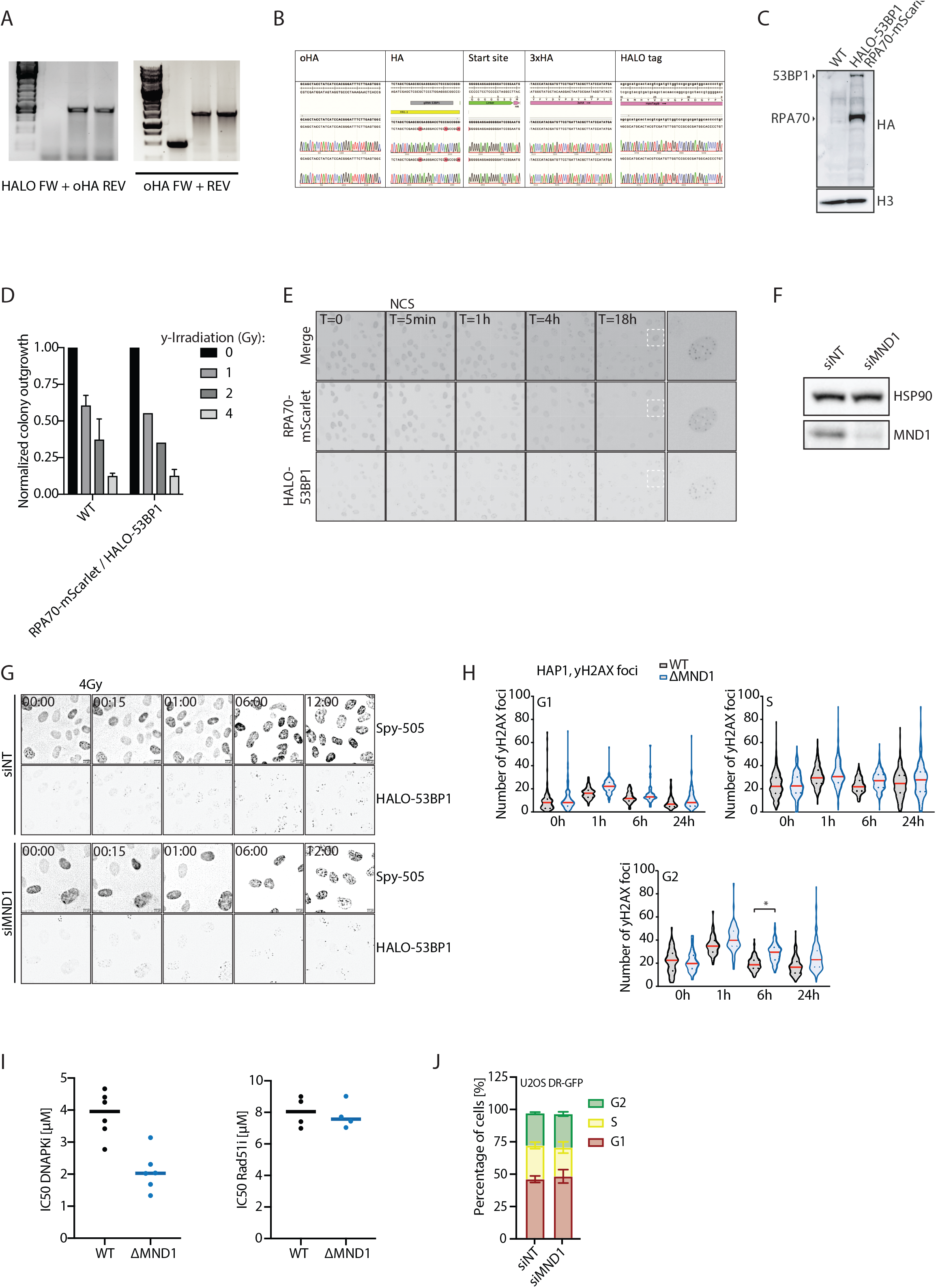
MND1 is involved in somatic HR. A)-E) Generation of RPE1 HALO-53BP1_RPA1-mScarlet2 cell line. A) Agarose gel depicting PCR amplification of inserted HALO- and HA-sequence at the 53BP1 locus. B) Sequencing result of inserted HALO- and HA-sequence at the 53BP1 locus. C) Western blot of WT and RPE1 HALO-53BP1_RPA1-mScarlet2 cell line with HA and histone H3 antibody. D) Clonogenic outgrowth of WT and RPE1 HALO-53BP1_RPA1-mScarlet2 cell line with increasing doses of IR. E) Example images of RPE1 HALO-53BP1_RPA1-mScarlet2 cell line after induction of DSBs with NCS. F) Western blot of siNT and siMND1 transfected RPE1 HALO-53BP1_RPA1-mScarlet2_GFP-MND1 cells. G) Example images of RPE1 HALO-53BP1_RPA1-mScarlet2 imaging with NT and MND1 siRNAs. DNA was stained using SPY-DNA 505. H) Immunofluorescence staining of HAP1 WT and ΔMND1 cells for yH2AX foci formation after 2Gy of IR. yH2AX foci number is depicted in G1, S and G2 phases separately. Three independent replicates are combined, and statistical analysis (t-test) was performed on mean values (* p ≤ 0.05). I) IC50 values of drug response assays for DNAPKi and RAD51i in HAP1 WT and ΔMND1 cells. J) Cell cycle profile of U2OS DR-GFP cells after treatment with siNT or siMND1.

**Suppl. figure 3:**
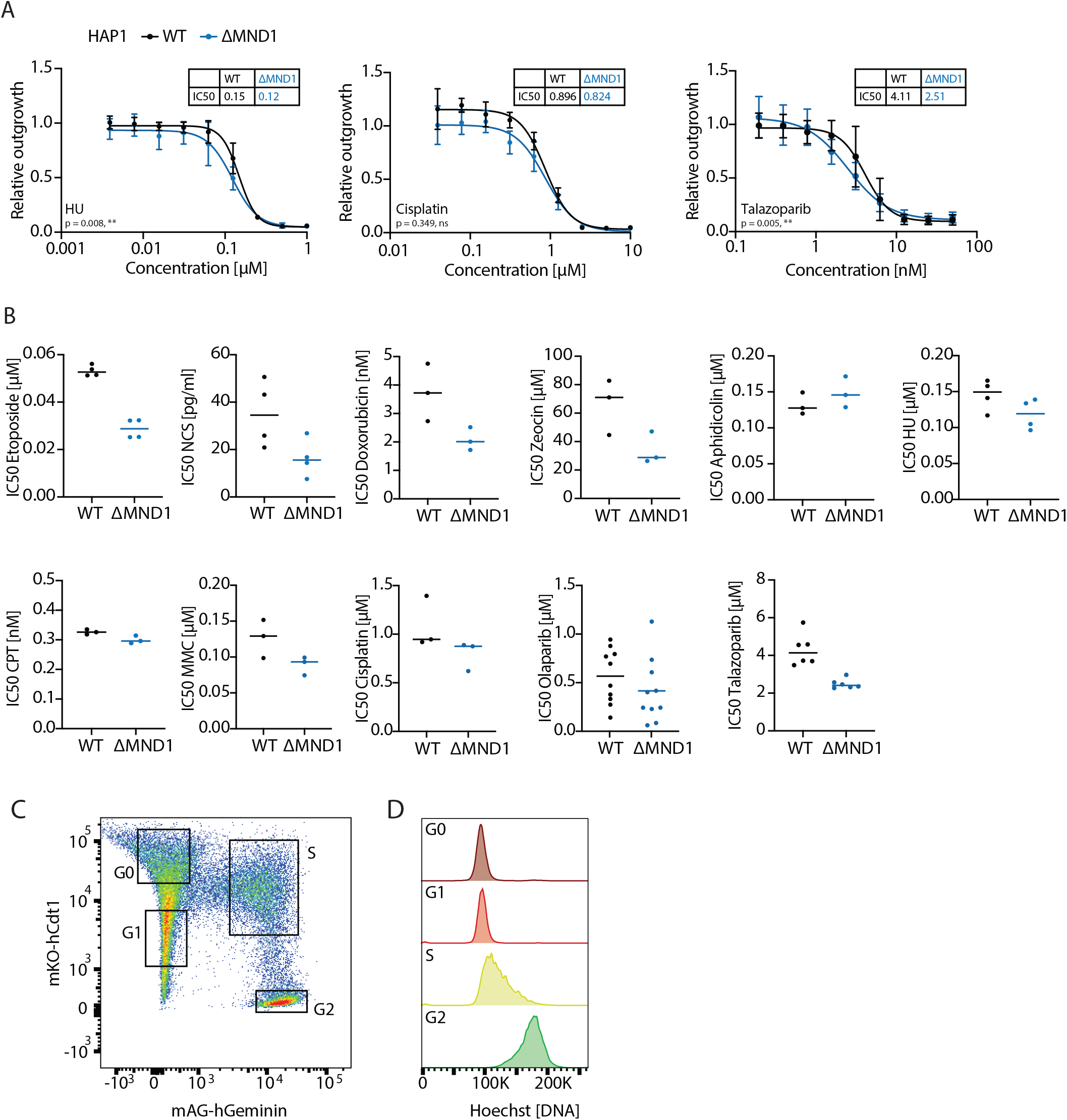
Loss of the MND1-HOP2 complex sensitizes cells towards DSB induction. A) Drug-response assay with hydroxyurea (HU, N=4), cisplatin (N=3) and talazoparib (N=6) in HAP1 WT and ΔMND1 cells. Data presented as mean±SD. Statistical analysis is performed on IC50 values and depicted in B). B) IC50 values of drug-response curves in HAP1 WT and ΔMND1 cells after treatment with etoposide, NCS, doxorubicin, zeocin, aphidicolin, HU, CPT, MMC, cisplatin, olaparib and talazoparib. C)&D) Flow cytometry analysis of Hoechst intensity in different sort gates of RPE1 Fucci cells depicted in Figure 3D.

**Suppl. figure 4:**
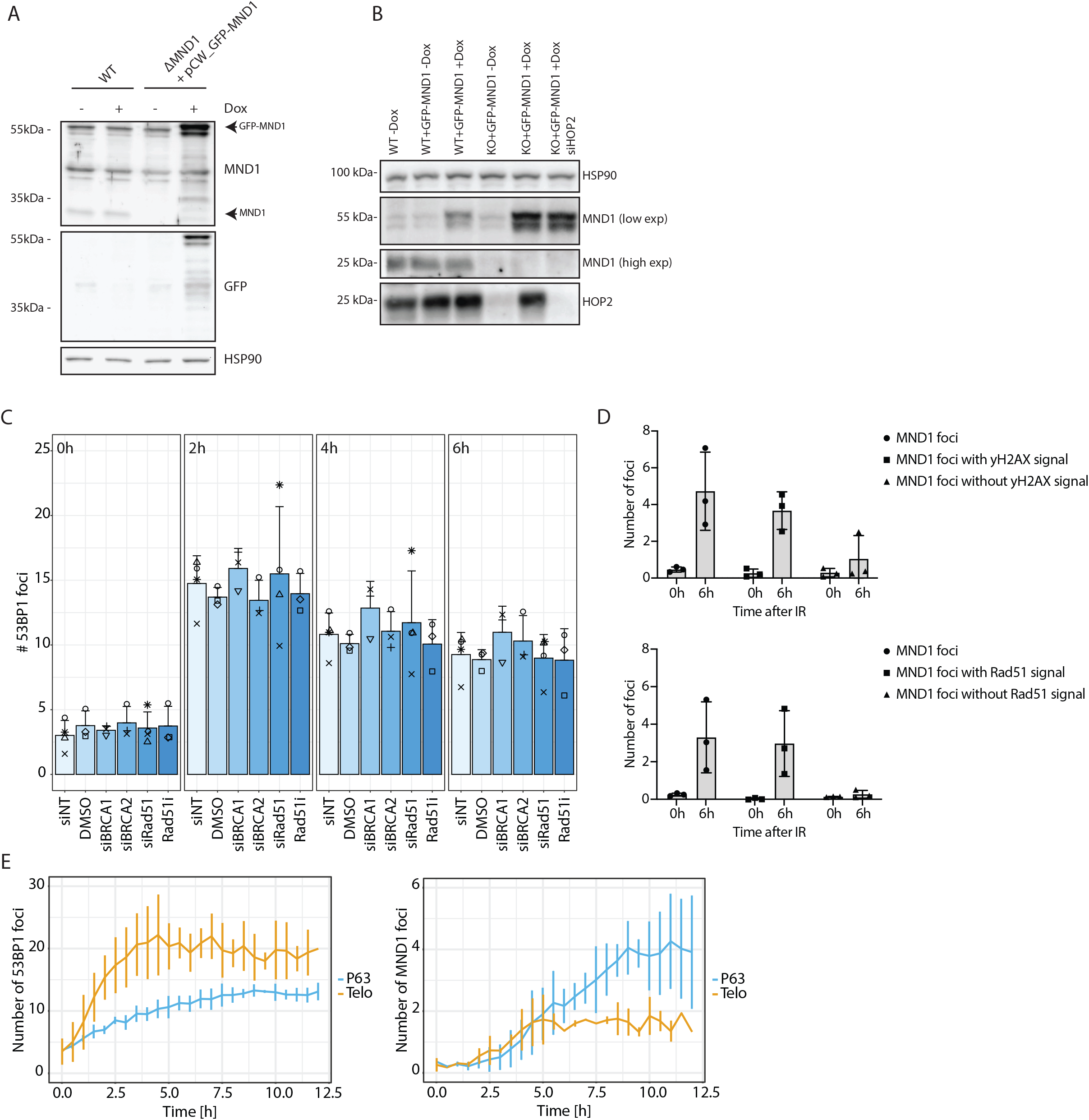
Foci formation of GFP-MND1 in RPE1 cells. A) Western blot of RPE1 HALO-53BP1_RPA1-mScarlet2 WT and ΔMND1 + GFP-MND1 cells. MND1 and GFP levels are depicted without Doxycycline and with Doxycycline addition. B) Western blot of RPE1 HALO-53BP1_RPA1-mScarlet2 WT and ΔMND1 + GFP-MND1 cells with or without doxycycline addition and siHOP2 treatment. Depicted are HSP90, MND1 and HOP2 levels. C) Number of 53BP1 foci in RPE1 HALO-53BP1, GFP-MND1 cells 0, 2, 4, and 6h after 4Gy of IR. D) MND1 foci numbers corresponding to Fig. 4 E) and F). E) Number of 53BP1 and MND1 foci and percentage of 53BP1 foci with MND1 in RPE1 dTag GFP-MND1 cells transfected with crRNAs targeting telomeres (Telo) or GAPDH pseudogenes (P63, (Berg et al., 2019)). Mean±SD is depicted of two independent replicates.

**Suppl. figure 5:**
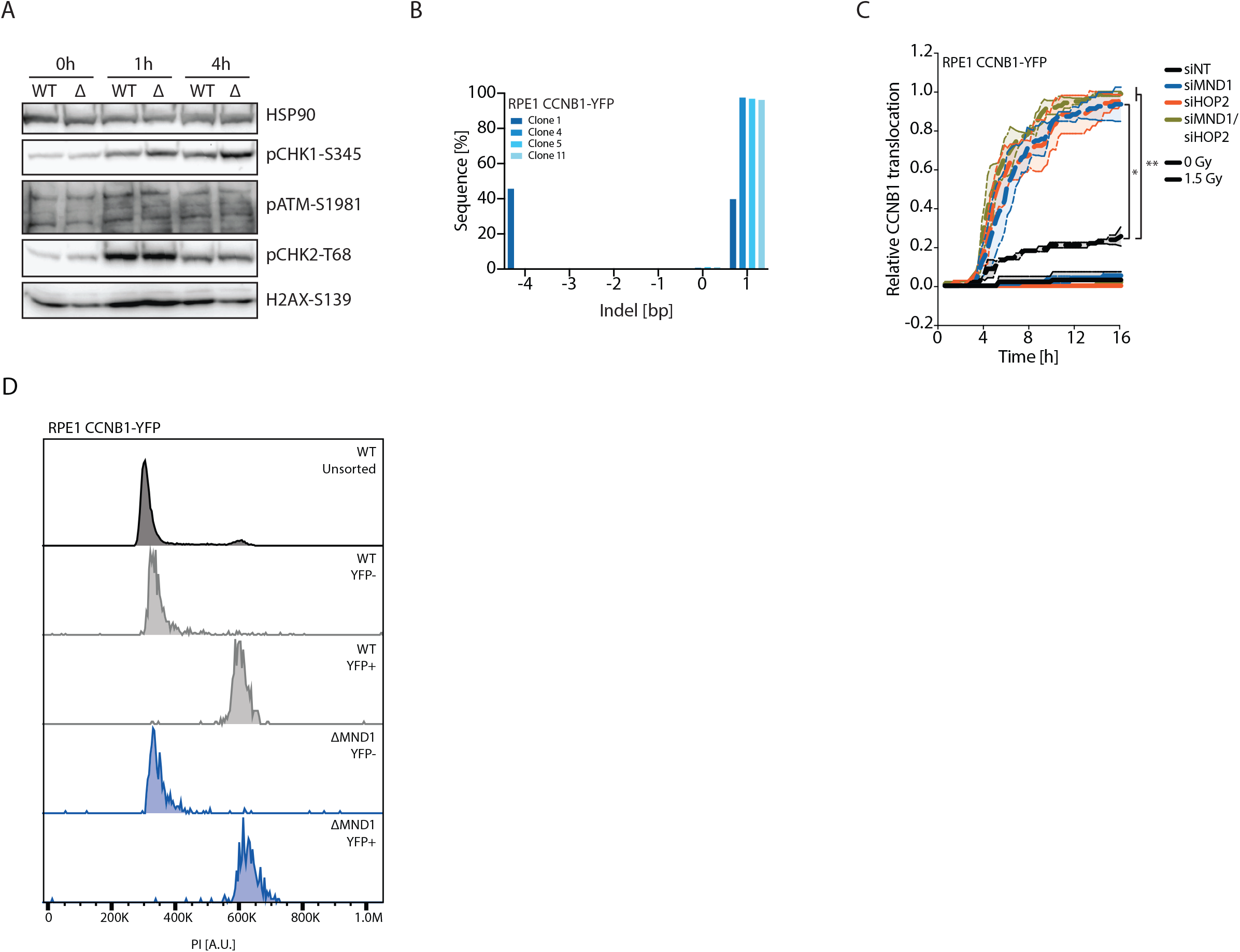
Checkpoint activation in MND1 deficient cells. A) Immunoblotting of checkpoint activation in HAP1 WT and ΔMND1 cells at 0h, 1h and 4h after 2Gy. B) Indel pattern of RPE1 CCNB1-YFP ΔMND1 clones. C) CCNB1 translocation after either 0Gy or 1.5Gy of IR in RPE1 CCNB1-YFP cells over time. Cells were treated with NT, MND1 and HOP2 siRNAs. Data is presented as mean±SD. D) DNA profile (PI) after FACS sorting of RPE1 CCNB1-YFP WT and ΔMND1 cells into YFP- and YFP+ populations.

## Notes

### Competing Interest Statement

The authors have declared no competing interest.

## References

Berg, J. van den et al. (2019) ‘DNA end-resection in highly accessible chromatin produces a toxic break’, bioRxiv. Cold Spring Harbor Laboratory, p. 691857. doi: 10.1101/691857.

Blackford, A. N. and Jackson, S. P. (2017) ‘ATM, ATR, and DNA-PK: The Trinity at the Heart of the DNA Damage Response’, Molecular Cell. Cell Press, 66(6), pp. 801–817. doi: 10.1016/J.MOLCEL.2017.05.015.

Blomen, V. A. et al. (2015) ‘Gene essentiality and synthetic lethality in haploid human cells.’, Science (New York, N.Y.). American Association for the Advancement of Science, 350(6264), pp. 1092–6. doi: 10.1126/science.aac7557.

Bonilla, B. et al. (2020) ‘RAD51 Gene Family Structure and Function Braulio’, Annu Rev Genet., 54, pp. 25–46. doi: 10.1146/annurev-genet-021920-092410.RAD51.

Brinkman, E. K. et al. (2014) ‘Easy quantitative assessment of genome editing by sequence trace decomposition’, Nucleic Acids Research. Oxford University Press, 42(22). doi: 10.1093/nar/gku936.

Bugreev, D. V. et al. (2014) ‘HOP2-MND1 modulates RAD51 binding to nucleotides and DNA’, Nature Communications. Nature Publishing Group, 5(1), p. 4198. doi: 10.1038/ncomms5198.

Chang, H. H. Y. et al. (2017) ‘Non-homologous DNA end joining and alternative pathways to double-strand break repair’, Nature Reviews Molecular Cell Biology. Nature Publishing Group, 18(8), pp. 495–506. doi: 10.1038/nrm.2017.48.

Chatterjee, N. and Walker, G. C. (2017) ‘Mechanisms of DNA damage, repair, and mutagenesis’, Environmental and Molecular Mutagenesis. John Wiley & Sons, Ltd, 58(5), pp. 235–263. doi: 10.1002/em.22087.

Chi, P. et al. (2007) ‘Bipartite stimulatory action of the Hop2-Mnd1 complex on the Rad51 recombinase’, Genes and Development, 21(14), pp. 1747–1757. doi: 10.1101/gad.1563007.

Cho, N. W. et al. (2014) ‘Interchromosomal homology searches drive directional ALT telomere movement and synapsis’, Cell. Elsevier Inc., 159(1), pp. 108–121. doi: 10.1016/j.cell.2014.08.030.

Ciccia, A. and Elledge, S. J. (2010) ‘The DNA Damage Response: Making It Safe to Play with Knives’, Molecular Cell. Cell Press, 40(2), pp. 179–204. doi: 10.1016/J.MOLCEL.2010.09.019.

Deans, A. J. and West, S. C. (2011) ‘DNA interstrand crosslink repair and cancer’, Nature Reviews Cancer. Nature Publishing Group, 11(7), pp. 467–480. doi: 10.1038/nrc3088.

Dev, H. et al. (2018) ‘and counters homologous recombination in’, Nature Cell Biology. Springer US, 20(August). doi: 10.1038/s41556-018-0140-1.

Dong, J. et al. (2018) ‘Inactivation of DNA-PK by knockdown DNA-PKcs or NU7441 impairs non-homologous end-joining of radiation-induced double strand break repair’, Oncology Reports, 39(3), pp. 912–920. doi: 10.3892/or.2018.6217.

Feringa, F. M. et al. (2016) ‘Hypersensitivity to DNA damage in antephase as a safeguard for genome stability’, Nature Communications. Nature Publishing Group, 7. doi: 10.1038/ncomms12618.

Feringa, F. M. et al. (2018) ‘Persistent repair intermediates induce senescence’, Nature Communications. Springer US, 9(1). doi: 10.1038/s41467-018-06308-9.

Francica, P. et al. (2020) ‘Functional Radiogenetic Profiling Implicates ERCC6L2 in Non-homologous End Joining ll ll Functional Radiogenetic Profiling Implicates ERCC6L2 in Non-homologous End Joining’, CellReports. The Authors, 32(8), p. 108068. doi: 10.1016/j.celrep.2020.108068.

Gerton, J. L. and Derisi, J. L. (2002) Mnd1p: An evolutionarily conserved protein required for meiotic recombination. Available at: www.pnas.orgcgidoi10.1073pnas.102167899.

Gunn, A. and Stark, J. M. (2012) ‘I-SceI-based assays to examine distinct repair outcomes of mammalian chromosomal double strand breaks’, Methods in Molecular Biology, 920, pp. 379–391. doi: 10.1007/978-1-61779-998-3_27.

Gupta, R. et al. (2018) ‘DNA Repair Network Analysis Reveals Shieldin as a Key Regulator of NHEJ and PARP Inhibitor Sensitivity’, Cell. Elsevier Inc., 173(4), pp. 972–988.e23. doi: 10.1016/j.cell.2018.03.050.

Hanahan, D. and Weinberg, R. A. (2011) ‘Hallmarks of cancer: The next generation’, Cell. Elsevier Inc., 144(5), pp. 646–674. doi: 10.1016/j.cell.2011.02.013.

Henry, J. M. et al. (2006) ‘Mnd1/Hop2 Facilitates Dmc1-Dependent Interhomolog Crossover Formation in Meiosis of Budding Yeast’, Molecular and Cellular Biology. American Society for Microbiology, 26(8), pp. 2913–2923. doi: 10.1128/mcb.26.8.2913-2923.2006.

Huang, F. and Mazin, A. V. (2014) ‘A small molecule inhibitor of human RAD51 potentiates breast cancer cell killing by therapeutic agents in mouse xenografts’, PLoS ONE, 9(6). doi: 10.1371/journal.pone.0100993.

Huang, R. X. and Zhou, P. K. (2020) ‘DNA damage response signaling pathways and targets for radiotherapy sensitization in cancer’, Signal Transduction and Targeted Therapy. Springer US, 5(1). doi: 10.1038/s41392-020-0150-x.

Hustedt, N. and Durocher, D. (2017) The control of DNA repair by the cell cycle, Nature Publishing Group. doi: 10.1038/ncb3452.

Jackson, S. P. and Bartek, J. (2009) ‘The DNA-damage response in human biology and disease’, Nature, pp. 1071–1078. doi: 10.1038/nature08467.

Jasin, M. and Rothstein, R. (2013) ‘Repair of strand breaks by homologous recombination’, Cold Spring Harbor Perspectives in Biology, 5(11). doi: 10.1101/cshperspect.a012740.

Jensen, R. B., Carreira, A. and Kowalczykowski, S. C. (2010) ‘Purified human BRCA2 stimulates RAD51-mediated recombination’, Nature. Nature Publishing Group, pp. 4–11. doi: 10.1038/nature09399.

Kelliher, J., Ghosal, G. and Leung, J. W. C. (2021) ‘New answers to the old RIDDLE: RNF168 and the DNA damage response pathway’, FEBS Journal, pp. 1–14. doi: 10.1111/febs.15857.

Krenning, L. et al. (2014) ‘Transient activation of p53 in G2 phase is sufficient to induce senescence’, Molecular Cell. Cell Press, 55(1), pp. 59–72. doi: 10.1016/j.molcel.2014.05.007.

Leu, J.-Y., Chua, P. and Roeder, S. (1998) ‘The Meiosis-Specific Hop2 Protein of S. cerevisiae Ensures Synapsis between Homologous Chromosomes’, Cell, 94, pp. 375–386.

Marshall, C. H. and Antonarakis, E. S. (2020) ‘Therapeutic targeting of the DNA damage response in prostate cancer’, Current Opinion in Oncology, 32(3), pp. 216–222. doi: 10.1097/CCO.0000000000000617.

Nickolo, J. A., Sharma, N. and Taylor, L. (2020) ‘Clustered DNA Double-Strand Breaks : Biological E ff ects and Relevance to Cancer Radiotherapy’.

Noordermeer, S. M. et al. (2018) ‘The shieldin complex mediates 53BP1-dependent DNA repair’, Nature. Nature Publishing Group, 560(7716), pp. 117–121. doi: 10.1038/s41586-018-0340-7.

O’Connor, M. J. (2015) ‘Targeting the DNA Damage Response in Cancer’, Molecular Cell. Cell Press, pp. 547–560. doi: 10.1016/j.molcel.2015.10.040.

Olivieri, M. et al. (2020) ‘A Genetic Map of the Response to DNA Damage in Human Cells’, Cell. Elsevier, 182(0), pp. 1–16. doi: 10.1016/J.CELL.2020.05.040.

Petukhova, G. V. et al. (2005) ‘The Hop2 and Mnd1 proteins act in concert with Rad51 and Dmc1 in meiotic recombination’, Nature Structural and Molecular Biology, 12(5), pp. 449–453. doi: 10.1038/nsmb923.

Petukhova, G. V., Romanienko, P. J. and Camerini-Otero, R. D. (2003) ‘The Hop2 protein has a direct role in promoting interhomolog interactions during mouse meiosis’, Developmental Cell, 5(6), pp. 927–936. doi: 10.1016/S1534-5807(03)00369-1.

Pezza, R. J. et al. (2007) ‘Hop2/Mnd1 acts on two critical steps in Dmc1-promoted homologous pairing’, Genes and Development, 21(14), pp. 1758–1766. doi: 10.1101/gad.1562907.

Pezza, R. J., Camerini-Otero, R. D. and Bianco, P. R. (2010) ‘Hop2-Mnd1 condenses DNA to stimulate the synapsis phase of DNA strand exchange’, Biophysical Journal. Biophysical Society, 99(11), pp. 3763–3772. doi: 10.1016/j.bpj.2010.10.028.

Pickett, H. A. and Reddel, R. R. (2015) ‘Molecular mechanisms of activity and derepression of alternative lengthening of telomeres’, Nature Structural and Molecular Biology. Nature Publishing Group, pp. 875–880. doi: 10.1038/nsmb.3106.

Pilié, P. G. et al. (2019) ‘State-of-the-art strategies for targeting the DNA damage response in cancer’, Nature Reviews Clinical Oncology. Nature Publishing Group, pp. 81–104. doi: 10.1038/s41571-018-0114-z.

Semlow, D. R. and Walter, J. C. (2021) ‘Mechanisms of Vertebrate DNA Interstrand Cross-Link Repair’, Annual Review of Biochemistry, 90, pp. 107–135. doi: 10.1146/annurev-biochem-080320-112510.

Shaltiel, I. A. et al. (2014) ‘Distinct phosphatases antagonize the p53 response in different phases of the cell cycle’, Proceedings of the National Academy of Sciences of the United States of America, 111(20), pp. 7313–7318. doi: 10.1073/pnas.1322021111.

Shaltiel, I. A. et al. (2015) ‘The same, only different – DNA damage checkpoints and their reversal throughout the cell cycle’, Journal of Cell Science. Company of Biologists Ltd, pp. 607–620. doi: 10.1242/jcs.163766.

Shiloh, Y. (2001) ‘ATM and ATR: Networking cellular responses to DNA damage’, Current Opinion in Genetics and Development, 11(1), pp. 71–77. doi: 10.1016/S0959-437X(00)00159-3.

Sobinoff, A. P. and Pickett, H. A. (2017) ‘Alternative Lengthening of Telomeres: DNA Repair Pathways Converge’, Trends in Genetics. Elsevier Ltd, pp. 921–932. doi: 10.1016/j.tig.2017.09.003.

Sullivan, M. R. and Bernstein, K. A. (2018) ‘RAD-ical New Insights into RAD51 Regulation’. doi: 10.3390/genes9120629.

Toulany, M. (2019) ‘Targeting DNA Double-Strand Break Repair Pathways to Improve Radiotherapy Response’, 1, pp. 1–20. doi: 10.3390/genes10010025.

Tsubouchi, H. et al. (2020) ‘Two auxiliary factors promote Dmc1-driven DNA strand exchange via stepwise mechanisms’, Proceedings of the National Academy of Sciences of the United States of America, 117(22). doi: 10.1073/pnas.1917419117.

Tsubouchi, H. and Roeder, G. S. (2002) ‘The Mnd1 Protein Forms a Complex with Hop2 To Promote Homologous Chromosome Pairing and Meiotic Double-Strand Break Repair’, Molecular and Cellular Biology, 22(9), pp. 3078–3088. doi: 10.1128/mcb.22.9.3078-3088.2002.

Tubbs, A. and Nussenzweig, A. (2017) ‘Endogenous DNA Damage as a Source of Genomic Instability in Cancer’, Cell, 168(4), pp. 644–656. doi: 10.1016/j.cell.2017.01.002.

Zhao, W. and Sung, P. (2015) ‘Significance of ligand interactions involving Hop2-Mnd1 and the RAD51 and DMC1 recombinases in homologous DNA repair and XX ovarian dysgenesis’, Nucleic Acids Research, 43(8), pp. 4055–4066. doi: 10.1093/nar/gkv259.

Zierhut, C. et al. (2004) ‘Mnd1 Is Required for Meiotic Interhomolog Repair’, Current Biology. Cell Press, 14(9), pp. 752–762. doi: 10.1016/J.CUB.2004.04.030.

